# Defective mitochondrial-lysosomal axis promotes extracellular vesicles release of mitochondrial components in Huntington’s Disease

**DOI:** 10.1101/2022.02.13.480262

**Authors:** Margarida Beatriz, Rita Vilaça, Sandra I. Anjo, Bruno Manadas, Cristina Januário, A. Cristina Rego, Carla Lopes

## Abstract

Mitochondrial and autophagy dysfunction are mechanisms proposed to be involved in the pathogenesis of several neurodegenerative diseases. Huntington’s disease (HD) is a progressive neurodegenerative disorder associated with mutant Huntingtin-induced abnormalities in neuronal mitochondrial dynamics and quality control. Former studies suggest that the removal of defective mitochondria may be compromised in HD. The mitochondrial quality control is a complex, well-orchestrated pathway that can be compromised through mitophagy dysregulation or impairment in the mitochondrial-lysosomal axis. Another mitochondrial stress response is the generation of mitochondrial-derived vesicles that fuse with the endolysosomal system and form multivesicular bodies that are extruded from cells as extracellular vesicles (EVs).

In this study, we comprehensively characterized the mitochondrial and autophagy alterations in premanifest and manifest HD patients and performed a proteomic and genomic EVs profile. We observed that manifest HD patients exhibit mitochondrial and autophagy impairment associated with enhanced EVs release. Further, we detected mitochondrial components in EVs released by HD cells and in neuron-derived EVs. The EV-associated mtDNA copies were elevated in manifest HD patients suggesting to be an alternative pathway for secretion of reactive mitochondrial components. This study provides a novel framework connecting EVs enhanced release of mitochondrial components to mitochondrial and lysosomal dysfunction in HD.

## INTRODUCTION

Huntington’s disease (HD) is an autosomal dominant neurodegenerative disease caused by an elongation of the CAG triplet in the exon 1 of the *huntingtin* gene (MacDonald et al., 1993). This pathology affects mainly the medium spiny neurons in the basal ganglia and to a lesser extent cortical areas that together drive motor coordination (Mehrabi et al., 2016; Vonsattel et al., 2008). Hence, involuntary movements are one of the symptoms jointly with early manifestations including cognitive and emotional alterations (Beglinger et al., 2010; Lemiere et al., 2004; Thompson et al., 2012). Although the monogenic nature of the disease has been recognized for almost three decades, there is no cure or treatment to mitigate the progression. The study of biomarkers specific for different stages of HD is a step forward to monitor patients’ conditions at the early stages of diagnosis and to adapt possible treatments. Until now the studies failed to show a consistently reproducible and clinically valid biomarker for HD, although numerous efforts are underway for the discovery of a reliable biomarker.

Mitochondrial dysfunction has been associated with neurodegenerative processes, including HD pathophysiology (reviewed in Carmo et al., 2018). In accordance, several mitochondrial parameters are altered in HD including ATP production deficit, increased reactive oxygen species (ROS) production, fragmented morphology and reduced mitochondrial movement (Lopes, 2020; Shirendeb et al., 2012). In line with these results, other studies have described the negative impact of mutant huntingtin (HTT) on mitophagy machinery, leading to dysfunctional mitochondria accumulation (Franco-Iborra et al., 2021). The mitochondrial protein quality control is tightly regulated by processes such as mitophagy, dynamics and molecular stress responses. When those mechanisms fail to counteract mitochondrial damages, there is an accumulation of dysfunctional mitochondria leading to enhanced oxidative stress, bioenergetic deficit and neurodegeneration (Jędrak et al., 2018; Lopes, 2022).

Extracellular vesicles (EVs) are nanosized particles (50–200 nm in diameter) that act as intercellular communicators. EVs are secreted by all cell types, including neurons, microglia and fibroblasts to transport cell-specific molecules like proteins, lipids, RNA, and DNA fragments from donor cells that can alter cell gene expression and function of recipient cells (O’Brien et al., 2020). The study of EVs dynamics has expanded exponentially over the past few years, particularly in neurodegenerative diseases. EVs cargo can be used as a diagnostic biomarker and as a pathology potentiator in the development and progression of neurodegenerative diseases (reviewed in Beatriz&Vilaça, 2021).

Recent work has revealed an intimate interconnection between classical autophagy machinery and autophagy-dependent EVs pathways that facilitates the disposal of unwanted genetic material and proteins (Leidal et al., 2020). Additionally, recent evidence suggests that EVs transfer mitochondrial DNA (mtDNA) and mitochondrial proteins intercellularly (Pasquale Sansone et al., 2017; Kiran Todkar et al., 2021; Xiaowan Wang et al., 2020); however, the biological relevance, the mechanisms underlying this process and the contribution in pathophysiological contexts are not clear. Neuronal-derived EVs (NDE) are a subpopulation of the central nervous system (CNS)-derived EVs that can be isolated from human fluids, including blood plasma and cerebrospinal fluid, to investigate their neuron-specific contents (Yousif et al., 2021). The presence of specific neuronal biomarkers in NDE isolated from blood samples of amyotrophic lateral sclerosis (ALS), Alzheimer and Parkinson disease patients, acknowledging their potential to monitor disease status and progression (Jia et al., 2020; Katsu et al., 2019; Niu et al., 2020).

With the objective of studying the crosstalk between the release of EVs and malfunction of dysfunctional mitochondria discard processes, in this work we characterized mitochondrial function and autophagy/mitophagy dynamics in cells from HD patients and controls (CTRs). Additionally, we isolated EVs secreted by these cells and NDE from the same patients’ biofluids and analyzed proteomic and genomic content. Our findings suggest that mitochondrial dysfunction associated with impaired autophagy enhances EVs release and this mechanism could act as an alternative pathway for the secretion of mitochondrial cargo in HD.

## MATERIALS AND METHODS

### Cell culture and reagents

Human dermal fibroblasts were obtained from skin explants of patients and CTRs, as described previously (Lopes, 2022). Fibroblasts were cultured in DMEM medium (Gibco), supplemented with 9 mM sodium bicarbonate, 10% fetal bovine serum (FBS) (Gibco) and 1% penicillin/streptomycin (Gibco). Cells were maintained in an incubator at 37 ^º^C with 5% CO_2_ for no more than 15 passages.

Dermal fibroblasts were successfully reprogrammed into induced-pluripotent stem cells (iPSC) using non-integrating Sendai virus for transduction, as described previously (Schlaeger et al., 2015). iPSC were grown and maintained in Geltrex® with StemFlex medium (Gibco) and the generation and proliferation of neural stem cells (NSC) was done as previously (Lopes, 2020). Briefly, to differentiate NSC toward a striatal-specific neuronal phenotype, until day five of differentiation, neuronal induction media was supplemented with 5 μM Dorsomorphin (Sigma) and 10 μM SB431542 (Peprotech) and later added 1 μM XAV 939 (Peprotech) and 200 ng/ml sonic hedgehog (SHH C-25II) (R&D) until day 12 (Chambers et al., 2009; Delli Carri et al., 2013; Nicoleau et al., 2013). At that point, NSC were dissociated with accutase (GRiSP), and plated on Geltrex®-coated coverslips in 24-well plates. To further obtain mature striatal-like neurons, from day 15 to 25 N2/B27 medium was supplemented with 200 ng/ml SHH, 1μM XAV939 and 30 ng/ml brain-derived neurotrophic factor (BDNF) (Peprotech). From day 26 onward only 50 ng/ml BDNF was added to the medium and medium changed every three days (Figure S1A.)

### Construction of plasmids and Viral production

The lentivirus pLESIP-GFP-LC3 was a kind gift from Dr. Ola Awad (University of Maryland, USA). pCL6EGwo and pCL6-CD63eGFP were a kind gift from Dr. Helmut Hanenberg and Dr. Bernd Giebel (University of Duisburg-Essen, Germany). The 86 bp insert of Mito-DsRed2 (Clontech) was subcloned at the NheI and BsrGI sites of the pCL6EGwo vector using standard recombinant DNA technology (pCL6-Mito-DsRed2). The pCL6-CD63eDsRed was constructed by cloning the 709 bp CD63 insert into the previous plasmid using NheI and *Bam*HI restriction. For the preparation of high-titer lentiviral stock lentiviral particles were produced in HEK293T cells by transiently co-transfecting the described plasmids with psPAX2 (#12260, Addgene) and pCMV-VSV-G (#8454, Addgene) using JetPrime reagent (Polyplus-transfection). After 48 hours, the viral particles were collected from the medium, purified and concentrated as described previously (Tiscornia et al., 2006). Lentiviral titration was estimated on FACS based on fluorescence.

### Incubation of fibroblasts with chloroquine and FCCP

With the objective of evaluating autophagic flux by western blotting, fibroblasts were incubated with 60 μM chloroquine (CQ) (Sigma) and 10 μM FCCP (CSNpharma) for 16 hours in EVs-depleted medium, using DMSO as control. EVs were isolated from the same cell culture media, as described. The number of EVs was assessed by Nanosight Tracking Analysis (NTA), and DNA was extracted to evaluate the mitochondrial DNA copies. As a control for EVs release, cells were incubated with 30 μM of GW4869 (CSNpharma) for 24 hours. For immunocytochemistry experiments, MitoDsRed and pLESIP-GFP-LC3 transduced cells were incubated with 100 μM CQ and 30 μM FCCP for five hours.

### Live cell microscopy

Fibroblasts were transduced either with pCL6-CD63eDsRed or pCL6-CD63eGFP vectors. Upon CD63+ expression, a 1:1 ratio was seeded for 24 hours in IBIDI μ-Dish, 35 mm high (IBIDI, Germany). Rapid time-lapse imaging was performed using a Zeiss Cell Observer Spinning Disk System (Carl Zeiss Microscopy) (63× oil1.4 NA PlanApochromat DIC objective) with an incubation system creating a 37 °*C* environment. Cells were imaged for up to six minutes. Particle tracking and trajectory analysis were performed with the Imaris x64 software version 9.5. Single-particle tracking trajectories are shown as dragon tail visualization. Speeds were derived from relative EVs displacement between two frames. The spots were quantified with a minimum xy diameter of 0.3 *μ*m, a max distance of 100 μm, the quality threshold of 600-3000 and a track duration of 1,5 s. The parameters analyzed were: Track length-total length of trajectory; Track straightness-relative measure for track straightness with 0 representing random diffusion and 1 representing a straight line.

### Cellular oxygen consumption and extracellular acidification rates

The oxidative phosphorylation and glycolytic levels of NSC were assessed through the measurement of the oxygen consumption rate (OCR) and extracellular acidification ratio (ECAR), respectively, on a Seahorse XFe96 apparatus (Agilent) as previously described (Lopes, 2020). One day before the experiments, NSC were seeded in a Geltrex®-coated XF96 Cell Culture Microplate at a density of 15 000 cells per well. For measuring the OCR, the apparatus added to cells, in the following order, 1 μM oligomycin (Alfa Aesar), 1 μM FCCP and 1 μM rotenone (Sigma) with 1 μM antimycin (Sigma). For measuring the ECAR, the apparatus added to the cell media, in the following order, 10 mM glucose (Sigma), 1 μM oligomycin (Alfa Aesar) and 50 mM 2-deoxyglucose (DOG) (Sigma). The final measurements were normalized to protein content using sulforhodamine B (Sigma) (Johnson et al., 2016).

### Transfection

NSC and neurons were transfected with Lipofectamine™ 3000 Transfection Reagent (Invitrogen™) according to the manufacturer’s protocol. Briefly, cells were plated in 24-well plates and transfected with 1.5 μg/μl of psDsRed-2Mito plasmid (Clontech).

### Mitochondrial movement

At day 80 of differentiation, transfected (psDsRed-2Mito) striatal-like neurons in coverslips were washed with Hanks’ Balanced Salt solution (HBSS) (#14025, Thermo Fisher Scientific). Mitochondrial movement videos were acquired on a Zeiss Cell Observer Spinning Disk System (Carl Zeiss Microscopy). For reference purposes, a static image of the neuron was acquired, and mitochondrial movement was recorded for three minutes with a 300 ms interval. Using the *TurboReg* plug on ImageJ, the video fluctuation was corrected. The *KymoToolBox* assembly of plugs produced a kymograph for each video, representing the mitochondrial movement across the timeline. Analyzed Kymo presented a set of parameters: velocity, persistence (Persistence=total distance traveled/distance in a straight line between the initial and final point; reflects the distance traveled in the same direction) distance traveled and time spent moving (anterograde-from the axon terminal to the soma-and retrograde-from the soma to the axon terminal movement) (Zala et al., 2013).

### Immunocytochemistry and image processing

Cells were washed with pre-warmed PBS at 37 °*C*, permeabilized with PHEM/0.1% Triton and fixed with PHEM/4% Paraformaldehyde (PFA) for 20 minutes. Blocking was performed for 40 minutes using PHEM/0.1% Triton/3% bovine serum albumin (BSA). The following primary antibodies were used: anti-P62 (SQSTM1) (1:100, #814802, BioLegend), anti-CD107b (LAMP-2) (1:100, #354301, BioLegend), anti-PINK1 (1:100, #846201, BioLegend), anti-LC3A/B (1:400, #12741S, Cell Signaling Technology), anti-OPA1 (1:50, #612606, BD Biosciences), anti-MFN2 (1:500, #M6319, Sigma), anti-DRP1 (1:100, #ab56788, Abcam), anti-nestin (1:200; #MAB1259 R&D), anti-gephyrin (1:200, #147111, Synaptic System), anti-synaptophysin (1:500, #ab32127, Abcam), anti-MAP2 (1:500, #ab92434, Abcam). After, cells were incubated with secondary antibodies for one hour at room temperature (RT) and mounted using Mowiol (Sigma); Hoechst (Invitrogen) was used to stain the nuclei. Images were acquired on a Zeiss LSM 710 Confocal System (Carl Zeiss Microscopy). Analysis of mitochondrial morphology was performed using Fiji ImageJ software. The area of the cell was established by drawing a ROI in the psDsRed-2Mito channel. The following mitochondrial parameters were obtained: number and area normalized to cell area; aspect ratio, which represents the elliptic shape of mitochondria; Feret’s diameter, which represents the distance between the two more far apart points of the mitochondria, reflecting the complexity of these organelles’ structure; Form factor that represents the branching of the mitochondrial net; and Circularity which describes more mature and elongated mitochondria for the value of 0 and circular and, so, more immature mitochondria for the value of 1 (Merrill et al., 2017). The equations that describe some of these parameters follow below: Aspect ratio= (major axis)/(minor axis); Form factor (perimeter)^2/(4π×area) and Circularity=(4π×area)/(perimeter)^2. Colocalization studies of fusion/fission proteins with mitochondria were performed on Z-stacks using the JACoP ImageJ plugin.

### Western blotting

Cells were lysed in a lysis buffer supplemented with 1:1000 protease inhibitor cocktail (Sigma), and protein was quantified using the Bio-Rad Protein Assay (Bio-Rad). For protein denaturation, SDS sample buffer was added to protein lysates (30 μg for fibroblasts, 25 μg for NSC) for five minutes at 95 °*C*. Samples were then loaded into 6-15% acrylamide gels. For EVs, the final isolated pellet (from 150 ml of fibroblasts or 60 ml of NSC culture media) was lysed in RIPA buffer and then protein was denatured in SDS sample buffer (non-reducing conditions were used to assess CD63 presence). Samples were then loaded into 10-12% acrylamide gels. Samples were subjected to SDS-PAGE run and transferred onto methanol-activated polyvinylidene difluoride (PVDF) Hybond-P membranes. Membranes were blocked in TBS-T/5% BSA for one hour at RT and incubated overnight at 4 °*C* with anti-P62 (SQSTM1) (1:500, #814802, BioLegend), anti-CD107b (LAMP-2) (1:250, #354301, BioLegend), anti-PINK1 (1:1000, #AP6406B, Abcepta), anti-LC3A/B (1:250) (#12741S, Cell Signaling), anti-β-Actin (1:5000, #D6A8, Cell Signaling) anti-β-Actin (1:5000, Sigma, #A5316), anti-OPA1 (1:500, #612606, BD Biosciences), anti-MFN2 (1:1000, #M6319, Sigma), anti-p-DRP (1:1000, #3455S, Cell Signaling), anti-DRP1 (1:1000, #ab56788, Abcam), anti-FIS1 (TTC11) (1:1000, #NB100-56646, Novus), anti-Alix (1:100, #sc-53538, Santa Cruz), anti-flotillin-2 (C42A3) (1:500, #3436,Cell Signaling), anti-calnexin (1:500) (#ab10286, Abcam) and anti-CD63 (TS63) (1:500, #10628D, Life Technologies). Primary antibodies were washed with TBS-T and membranes were incubated with secondary antibodies anti-mouse and anti-rabbit (1:5000) diluted in TBS-T/1% BSA for one hour at RT. Secondary antibodies were washed and immunoreactive bands were developed for five minutes with ECF (GE Healthcare) and revealed in a Chemidoc Imaging System (Bio-Rad) (Lopes, 2020).

### Dot blotting

Samples were loaded directly onto methanol-activated PVDF membranes and left to dry at RT. Membranes were blocked in TBS-T/5% BSA for one hour at RT and incubated overnight at 4 °*C* with anti-CD63 (TS63) (1:500, #10628D, Life Technologies), anti-CD171 (L1CAM) (1:200, #13-1719-82, Thermo Fisher) and anti-synaptophysin (YE269) (1:500, #ab32127, Abcam). The following procedure was similar to western blotting.

### EVs and NDE isolation and characterization

For EVs isolation, fibroblasts were cultured in EVs-depleted medium: FBS was centrifuged at 100 000 × *g* for 18 hours at 4 °*C* to pellet animal vesicles. For isolation of EVs of fibroblasts and NSC, cell culture supernatant was collected every other day and stored at -80 °*C* for posterior use. For isolation of EVs in bulk, culture media was thawed at 37 °*C* and filtered (0.22 μm) to remove cell debris and proceed as described (Théry et al., 2006). Briefly, medium was centrifuged at 10 000 × *g* for 30 minutes, and twice at 100 000 × *g* for 70 minutes to pellet EVs. EVs were resuspended in different buffers depending on the final use: PBS for DNA extraction, NTA and mass spectrometry; PBS/2% PAF for transmission electron microscopy (TEM) and RIPA for western/dot blotting.

### Isolation of EVs from blood plasma

Blood samples were obtained from HD patients and healthy CTRs upon informed consent. Blood was collected into collection tubes with K2EDTA (BD Vacutainer Plastic Blood Collection Tubes). Briefly, blood was diluted 1:1 in PBS and isolated by a Ficoll (GE Healthcare, Fisher Scientific) density gradient separation performed at 800 g for 30 minutes at RT, no break. Plasma was kept at -80 °*C* until further use.

EVs from blood plasma were isolated using the exoEasy Maxi kit (Qiagen) following the manufacturer’s protocol. The pellet was resuspended in 300 μl of Resuspension Buffer. The total plasma exosomes were stored at −80 °*C* to NTA and dot blot analysis and further isolation of NDE.

### Neural-enriched EV extraction

NDE were isolated from total EVs of plasma using the Basic Exo-Flow Capture Kit (System Biosciences) as described (Winston et al., 2019). First, 40 μl of streptavidin magnetic beads were incubated with 4 μg of anti-CD171 (L1CAM, neural adhesion protein) biotinylated antibody (clone 5G3, #13-1719-82, Thermo Fisher) for three hours on ice. After the beads were suspended in Bead Wash Buffer and 100 μl of total EVs previously isolated from plasma was added. To allow the magnetic beads to capture the desired vesicles, the complexes were left overnight gently rotating at 4 °*C*. The beads were washed and EVs/NDE were eluted in exosome elution buffer and stored at -80 °*C* until further required.

### Particle size and concentration analysis

EVs were resuspended in 1 ml of water containing low-mineral concentration. NTA analysis was performed in a NanoSight NS300 instrument with a sCMOS camera module (Malvern Panalytical), following general recommendations. Analysis settings were optimized and kept constant between samples. The videos were analyzed to obtain the mean size and estimated concentration of particles. Data were processed using the NTA 3.1 analytical software.

### TEM and immunogold-TEM

EVs were isolated from culture media of fibroblasts and NSC. The antibody anti-TFAM (1:150, #sc-28200, Santa Cruz) were used for immunoelectron labeling. Briefly, exosome samples were placed on carbon-Formvar coated 300 mesh nickel grids, and samples were allowed to absorb for 20 min. Following washing in PBS (2x 3 minutes), grids were floated (sample side down) onto 50 μl drops of 50 mM glycine (4x 3 minutes) to quench free aldehyde groups and then transferred to blocking solution (5% BSA, 0.01% saponin in PBS) for 10 minutes. Then, grids were incubated in blocking buffer (1% BSA, 0.01% saponin in PBS; negative control) or primary antibodies for two hours followed by washing steps in BSA 1% (6x 3 minutes). Secondary antibodies (anti-rabbit IgC immunogold conjugates, 1:200) were then added for one hour, followed by additional washing steps (8x 2 minutes). After, samples were fixed in glutaraldehyde 1% for five minutes and washed in water (8x 2 minutes). A final contrasting step was performed using uranyl-oxalate solution, pH 7, for five minutes followed by a mixture of 4% uranyl acetate and 2% methyl cellulose, for 10 minutes on ice. Imaging was conducted using a FEI-Tecnai G2 Spirit Bio Twin TEM at 100 kV.

### SWATH-MS analysis

The pellets of EVs were dissolved in 2x SDS-PAGE Sample Loading Buffer (Nzytech) aided by ultrasonication. Each sample was spiked with the same amount of a recombinant protein (MBP-GFP) to be used as internal standard (Anjo et al., 2019) and denatured at 95 °*C* for five minutes. A small part of each denatured sample was pooled to generate the proteome library (protein composition of each condition), and the remaining sample was used separately for SWATH-MS analysis to extract quantitative information. Samples were then alkylated with acrylamide and subjected to in-gel digestion by using the short-GeLC approach (Anjo et al., 2015).

All samples were analyzed on a NanoLC™ 425 System coupled to a Triple TOF™ 5600 mass spectrometer (Sciex®) using two acquisition modes: i) the pooled samples were analyzed by information-dependent acquisition (IDA) and, ii) the individual samples by SWATH-MS mode. The ionization source was the DuoSpray™ Ion Source (Sciex®) with a 50 μm internal diameter (ID) stainless steel emitter (NewObjective).

For IDA experiments, the mass spectrometer was set to scanning full spectra (m/z 350-1250) for 250 ms, followed by up to 100 MS/MS scans (*m/z* 100–1500) per cycle, in order to maintain a cycle time of 3.309s. The accumulation time of each MS/MS scan was adjusted in accordance with the precursor intensity (minimum of 30 ms for precursor above the intensity threshold of 1000). Candidate ions with a charge state between +2 and +5 and counts above a minimum threshold of 10 counts per second were isolated for fragmentation and one MS/MS spectrum was collected before adding those ions to the exclusion list for 25 s (mass spectrometer operated by Analyst® TF 1.7, Sciex). The rolling collision was used with a collision energy spread (CES) of 5. For SWATH experiments, the mass spectrometer was operated in a looped production mode (Gillet et al., 2012) and specifically tuned to a set of 60 overlapping windows, covering the precursor mass range of 350–1,250 m/z. A 200 ms survey scan (350–1,250 m/z) was acquired at the beginning of each cycle, and SWATH-MS/MS spectra were collected from 100 to 1,500 m/z for 50 ms resulting in a cycle time of 3.254 s. The collision energy for each window was determined according to the calculation for a charge +2 ion centered upon the window with variable CES, according to the window.

Peptide identification and library generation were performed by searching all the IDA samples using the ProteinPilot™ software (v5.1, ABSciex®) with the following parameters: i) search against a database from SwissProt composed by Homo Sapiens (release at March 2019) and MBP-GFP (IS) sequences; ii) acrylamide alkylated cysteines as fixed modification; and iii) trypsin as digestion type. An independent False Discovery Rate (FDR) analysis using the target-decoy approach provided with Protein Pilot software was used to assess the quality of the identifications and positive identifications were considered when identified proteins and peptides reached a 5% local FDR (Sennels et al., 2009; Tang et al., 2008). Complementary/Further information can be found in Supplementary material.

### Systematic analysis

Biological and molecular processes terms from Gene Ontology were analyzed using PANTHER (http://www.pantherdb.org/). The protein–protein interaction network and identification count were obtained from the STRING database (http://string-db.org/) and the EVpedia database (https://evpedia.info) respectively.

### DNA/RNA Extraction, cDNA and PCR

Genomic DNA was extracted using PureLink® Genomic DNA Kit and RNA with the PureZOL® RNA Isolation Reagent according to the manufacturer’s instructions. Reverse transcription was performed with iScript™ cDNA Synthesis Kit. Real-time PCR (qPCR) was performed with iQTM SYBR® Green Supermix on a CFX96 Touch™ Real-Time PCR Detection System as previously described (Lopes 2020). QPCR was performed according to the manufacturer’s protocol. 18S was used for internal reference gene. Expression values were calculated using the 2^−ΔΔCt^ method. All PCR samples were run in technical triplicates, and the average Ct-values were used for calculations. CAG repeats number in cells was determined by PCR, as described previously (Evers et al., 2015). Primers used are listed in Supplementary Table 1.

### Analysis of Mitochondrial DNA Copy Number

EVs were isolated from FBS Exo-depleted culture media of fibroblasts and NSC and NDEs were extracted from blood plasma. To eliminate contaminant DNA, EVs resuspended in PBS, were pretreated with 10 μg/μl of DNAse I (Canvax), according to the manufacturer’s instructions. EVs DNA (exoDNA) and cellular DNA were extracted with QIAamp DNA Micro Kit (QIAGEN) or DNeasy tissue and blood kit (QIAGEN) and DNA eluted in 20 μL DEPC water. The concentration and purity of DNA derived from exosomes was assessed by measurement of absorbance and 260/280 ratio with a NanoDrop 2000/c (Thermo Fisher Scientific).

To detect the presence of mtDNA in exosomes and cells, exoDNA (8 ng) or cell’s DNA (6 ng) was used for whole mtDNA amplification according to (Dames et al., 2015). Briefly, long-range PCR was used to amplify the whole mtDNA sequence using three overlapping primer pairs. Each amplification was performed using Phusion® High-Fidelity DNA polymerase (Thermo Scientific) and the amplicons were separated on a 0,8% agarose gel and visualized with a GelDoc system (Bio-Rad). Primers used are listed in Supplementary Table 1.

### Quantification of mtDNA copies number

Quantification of absolute mtDNA copy number was performed as described in (Gonzalez-Hunt et al., 2016), with minor modifications. The method is based on real-time PCR assay using a standard curve with amplifying serial dilutions of a mtDNA-specific gene. The average Ct value of each DNA sample was used to interpolate with the standard curve to calculate the absolute number of copies of mtDNA gene. The standard curve was generated by cloning a short 100-bp sequence of mitochondrial 16S rRNA gene into pGEM®-T Easy Vector (Promega). The 16S_rRNA gene sequence was amplified with Taq Polymerase (Supreme NZYtech II master mix) and cloned into pGEM-T easy vector. Briefly, 2 ng of ExoDNA was used to calculate mtDNA copies number with iQ™ SYBR® Green Supermix (Bio-Rad) on a CFX96 Touch™ Real-Time PCR Detection System (Bio-Rad). Logarithmic regression was used to plot Ct values against a known number of copies and absolute mtDNA copies number for each sample was calculated as previously described (Gonzalez-Hunt et al., 2016). Primers used are listed in Supplementary Table 1.

### Measurement of mitochondrial hydrogen peroxide production

Cells were plated in Geltrex®-coated 96-well plates at a density of 15 000 cells per well the day before the experiments. To study mitochondrial reactive oxygen species, cell media was removed, cells were washed with warm HBSS and incubated with 10 μM MitoPY1 (Sigma) for 20 minutes at 37 °*C*. Cells were washed twice with HBSS and MitoPY1 fluorescence (excitation-503 nm, emission - 530 nm) was measured using a Fluorimeter SpectraMax GEMINI EM (Molecular Devices) or a Microplate Reader SpectraMax iD3 (Molecular Devices). Hydrogen peroxide basal production was measured for 10 minutes and for 30 minutes after the addition of 3 μM myxothiazol or 1 μM rotenone. Results obtained reflect RFUs from myxothiazol/rotenone stimulus normalized to RFUs from control wells (Lopes, 2022).

### Statistical analysis

Statistical computations were performed using GraphPad Prism version 9.0, GraphPad Software, La Jolla, CA, USA, and SPSS version 21.0 (IBM SPSS Statistics for Windows, IBM Corp). Statistical analysis was performed for individual subjects and among groups (controls, pre-symptomatic and stage 3 symptomatic patients). Results are the mean±SEM of the indicated number of independent experiments in figures legends. For cell experiments, at least three independent assays were performed for each experimental condition. Statistical significance was analyzed using parametric test, one-way ANOVA, followed by Bonferroni post hoc test and non-parametric test Kruskal Wallis followed by Dunn’s multiple comparison test. Correlations were done using the Spearman rank correlation coefficient (ρ). P<0.05 was considered significant.

## RESULTS

### Mitochondrial function and dynamics in differentiated neuronal cells

We have previously demonstrated that human neural stem cells (NSC) from Huntington’s disease (HD) patients exhibit features of mitochondrial and metabolic dysfunction (Lopes, 2020). IPSC reprogrammed from patients’ fibroblasts (Lopes, 2022) were differentiated into progenitors and fully mature GABAergic striatal-like neurons using published protocols (Figure S1a) (Delli Carri et al., 2013; Lopes, 2020). The CAG repeat expansion was confirmed in HD cell lines by PCR (Figure 1a). The successful differentiation of the iPSC into NSC and GABAergic neurons was confirmed using cell-specific markers: Nestin for neural progenitors and gephyrin, synaptophysin and MAP2 to identify inhibitory synapses, respectively (Figure 1b).

**Figure 1.**
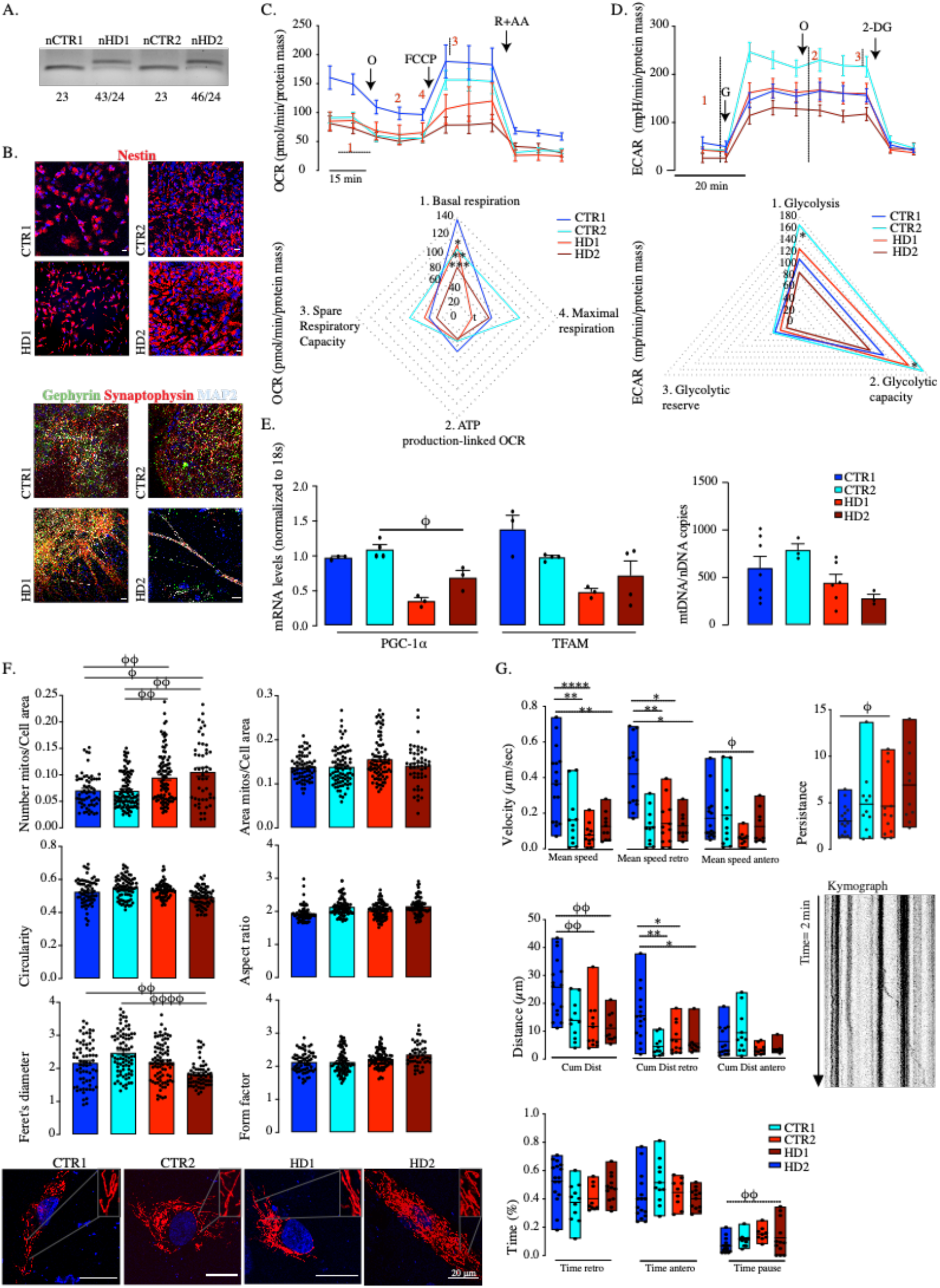
Mitochondrial bioenergetic and dynamic abnormalities in HD and control differentiated NSC and striatal-like neurons. (a) PCR analysis of the CAG repeat expansion. (b) Expression of neural progenitors’ marker Nestin at day 15 and colocalization of synaptic markers for inhibitory (synaptophysin with gephyrin) showing synapse formation in differentiated neurons (MAP2+) at day 80. (c) Cellular oxygen consumption rate (OCR) graph shown on top and spider plots on the bottom: (1) basal respiration; (2) oxygen consumed for ATP generation through the complex V; (3) Spare respiratory capacity; (4) maximal respiration capacity; (b) and extracellular acidification rate (ECAR) graph shown on top and spider plots on the bottom: (1) Glycolysis; (2) Glycolytic capacity, (3) Glycolytic reserve in NSC from HD patients and controls (N=5-10). Values for mean±S.E.M of at least 3 independent experiments. (e) mRNA levels of PGC-1α and TFAM normalized to 18S, and mitochondrial DNA (mtDNA) copy number (N=3-6). (f) Mitochondrial morphology, size and number in HD NSC vs CTR and representative images of mitochondria fluorescence after transfection with psDsRed-2Mito (scale bar=20 μm) (N=48-79). (g) Analysis of mitochondrial motion in striatal-like neurons transfected with psDsRed-2Mito and representative kymograph (N=10-15). Bar plots represent mean±S.E.M and floating bars show the minimum and maximum (line at mean). Statistical analysis: one-way ANOVA followed by Bonferroni post-hoc: * p< 0.05, ** p<0.01, *** p< 0.001, **** p<0.0001 or Kruskal-Wallis comparisons: ϕ p<0.05, ϕϕ p<0.01, ϕϕϕϕ p<0.0001.

First, we analyzed mitochondrial bioenergetics and dynamic events in neuronal progenitors. HD-NSC exhibited reduced levels of basal and maximal respiration compared to CTRs, and a tendency for lower spare respiratory capacity (Figure 1c, S1b). In addition, HD-NSC presented a reduction of glycolysis and glycolytic capacity (Figure 1d, Figure S1c). Together, these data suggest that HD neural progenitors have a decreased bioenergetic profile. Despite the metabolic alterations found in HD-NSC we did not find alterations in mitochondrial hydrogen peroxide production (Figure S1d, e).

The number of mtDNA copies has a crucial role in mitochondria biogenesis and bioenergetics, which is regulated by the nuclear transcriptional factors PGC-1α (peroxisome proliferator-activated receptor gamma co-activator) and TFAM (mitochondrial transcription factor A). In accordance, analysis of the mRNA transcripts showed a significant decrease in the levels of PGC-1α and a tendency for lower mRNA TFAM in HD-NSC (Figure 1e). The quantification of mtDNA copies showed a trend for lower values in these cells despite absent statistical significance (Figure 1e).

Since mitochondria are organelles that depend highly on dynamic mechanisms to maintain quality control, we sought to evaluate mitochondrial fission/fusion processes. The total levels of fission proteins (phospho-Drp, Drp1 and FIS1) and fusion proteins (Mitofusin2 and OPA1) were unaltered between HD-NSC and CTRs, although Mitofusin2 and Drp1 colocalize less with mitochondria in HD-cells (Figure S2a, b). Mitochondrial morphological changes were further investigated in NSC transduced with pDsRed2Mito vector. A higher mitochondrial copy number in HD-NSC was associated with a significant decrease in the Feret’s diameter. The other mitochondrial shape descriptors were indistinguishable between mitochondrial subpopulations (Figure 1f). To gain further insight into mitochondrial dynamics, mitochondrial movement was evaluated in differentiated post-mitotic striatal-like neurons in which mitochondria are expected to be more mature than in proliferative NSC (Beckervordersandforth et al., 2017). We observed slower velocity and reduced distance for mitochondria in HD-cells associated with higher persistence meaning that they changed fewer times the direction of movement (Figure 1g).

### Auto(mito)phagy analysis in human fibroblasts

Our previous work described altered mitochondrial function and redox deregulation in fibroblasts from manifest HD patients (Lopes, 2022). Since dysfunctional/damaged mitochondria are removed through selective autophagy, more precisely mitophagy, we further analyzed the levels of proteins involved in this process in three premanifest HD (fpHD), two manifest HD (fHD) lines of fibroblasts (vs two CTRs) to infer autophagy at different disease stages of the disease (Figure 2). Since defects in autophagic flux have been linked to HD (Pircs et al., 2018), we monitored autophagic activity by evaluating the levels of LC3II, LC3-II/ I ratio, p62 (proteins related to autophagosomal assembly and selective autophagy), LAMP2 (lysosomal constitution) and PINK1 (mitophagy ubiquitin-dependent pathway) by western blot and immunocytochemistry. We found increased expression levels of the autophagic markers p62, LC3-II and LC3-II/LC3-I ratio as well as the mitophagy marker PINK1 in manifest HD fibroblasts lines (Figure 2a).

**Figure 2.**
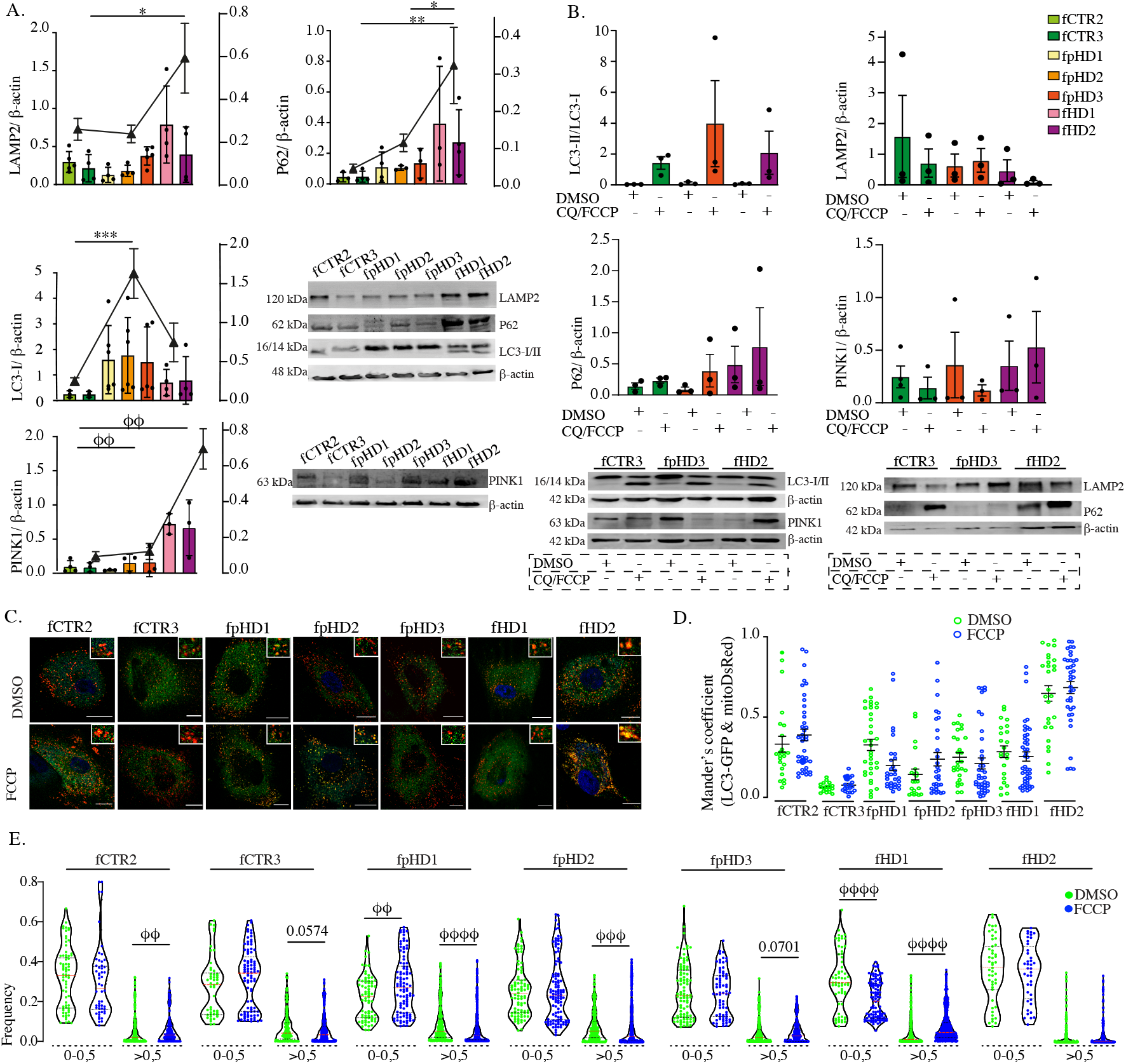
Basal and dynamic autophagic flux in fibroblasts from HD patients and controls. (a) Western blot analysis of autophagy-associated protein markers LAMP2, p62, LC3 and PINK1 in fibroblasts from HD patients and controls (2 control, 3 premanifest and 2 manifest HD). Fibroblasts were grouped for purposes of comparison as presented in the right YY axis. Results were normalized to ß-actin levels (N=4). (b) Analysis of autophagic flux by western blotting in one cell line per group treated with chloroquine (CQ) (60 μM) and FCCP (10 μM) for 24h to block lysosomal acidification and promote mitophagy. Results were normalized to ß-actin levels (N=3). (c) Colocalization represented by Mander’s coefficient, of LC3 and mitochondria in fibroblasts transduced with mitoDsRed and GFP-LC3 and treated with FCCP (30 μM) for 5h (N>=17) and representative images (scale bar=20 μm). (d) Size of LC3-puncta in fibroblasts treated with FCCP (30 μM) for 5h. Results were normalized to the total number of LC3-puncta. Area is represented in μm^2^ (XX axis) (N>=20). Bar plots represent mean±S.E.M and violin plots represent median with interquartile range. Statistical analysis: one-way ANOVA followed by Bonferroni multiple comparisons test: * p< 0.05, ** p<0.01, *** p< 0.001 or Kruskal-Wallis comparisons: ϕϕ p<0.01, ϕϕϕ p< 0.001, ϕϕϕϕ p<0.0001.

Only the cytosolic isoform of LC3 (LC3-I) was detected at basal levels in CTRs and fpHD. The lipidated form of LC3-I (LC3-II) is recruited to autophagosomal membranes and rapidly degraded in active lysosomes which is consistent with functional autophagy machinery in CTRs and pHD. (Figure 2a).

Additionally, we evaluated the autophagic flux in one fibroblast line from each group treated with CQ, which blocks protein degradation by lysosomal acidification, combined with FCCP to induce mitochondrial depolarization and fragmentation which precedes mitophagy. As expected, incubation with CQ/FCCP resulted in the accumulation of LC3II, an increase of LC3-II/LC3-I ratio and accumulated p62 protein particularly in pHD and HD cell lines (Figure 2b). Interestingly, the expression levels of PINK1 protein levels were only increased in manifest HD. Protein LAMP2 is essential in controlling autolysosome formation and degradation of autophagosomes (Abokyi et al., 2021). The levels of LAMP2 were decreased in manifest HD and control cells after treatment with the autophagy inhibitor CQ indicating downregulation of the lysosomal membrane-stabilizing and loss of lysosomal membrane integrity and function at a greater level than in pHD (Figure 2b).

Together, these data showed an accumulation of proteins involved in detecting, targeting and eliminating dysfunctional mitochondria suggesting impaired autophagosome-lysosome fusion or inhibition of lysosome degradative process in manifest HD.

We further investigated alterations in mitophagy through colocalization analysis in fibroblasts expressing a lysosomal fluorescence tagged protein (LC3-GFP) and a mitochondrial fluorescence tagged protein (mitoDsRed). We sought to evaluate the number, size, and colocalization of GFP-LC3 puncta with mitochondria under basal conditions and in the presence of FCCP to induce mitophagy or CQ to inhibit lysosomal degradation. The colocalization analysis of LC3 puncta with mitochondria showed no differences between treated and non-treated cell lines with FCCP, except for the one manifest HD cell line that showed an increased accumulation of LC3 puncta in mitochondria (Figures 2c,d). Interestingly, the pHD2 line also expressed more p62 and PINK1 proteins upon exposure to auto/mitophagy modulators. Impairment of lysosomal degradation resulted in an augmented area of LC3-positive structures (Figure S3a). The puncta area was distributed by categories assuming that normal sized had under 0.5 μm^2^ and enlarged more than 0.5 μm^2^. The increase in the area of LC3 puncta is cell-specific as we observed an enlargement in puncta in lines fCTR2, fpHD3 and in both manifest HD lines after acute mitophagy induction with FCCP (Figure 2d), suggesting an enlargement of the autophagosomes in HD-fibroblasts upon promotion of mitochondrial elimination. No changes or slight reduction in size was observed in fCTR3, fpHD1 and fpHD2 (Figure 2d). The number of LC3 puncta was also not altered as well as the total number of puncta per cell area. The histograms with the distribution of puncta in each area interval are represented for every fibroblast line treated whether with FCCP or CQ in supplementary data (Figure S3b, c). These data demonstrate that mitochondria seem to be more targeted for mitophagy in manifest HD cell lines.

### Fibroblasts and NSC from HD patients release more extracellular vesicles

To evaluate the release of small EVs (sEVs) by HD-fibroblasts, -NSC and CTRs lines we performed sequential ultracentrifugation to isolate these vesicles from the conditioned growth media (Figure 3a,b). Characterization of EVs population according to ISEV guidelines (Théry et al., 2018) included the presence of typical protein markers for sEVs such as Alix, Flotillin-2 and CD63, and calnexin was used as a negative control to exclude cellular contaminants (Figure 3c, f). The NTA for particle size showed that EVs from fibroblasts and NSC were in the expected diameter ranging from 50–200 nm (Figure 3d, g). In addition, the EVs released by these cells exhibited the typical cap-shaped morphology (Figure 3e, h). To further assess the EVs proteomic cargo, isolated EVs from HD- and CTR-NSC were analyzed by liquid chromatography-tandem mass spectrometry (LC-MS/MS) (Figure 3i, S4). A combined analysis of the replicates identified 82 common proteins between HD-NSC and CTRs. These cell type-specific proteins were further submitted for gene enrichment analysis including Gene Ontology Biological Processes (Figure 3i). The analysis revealed an enrichment of proteins related to vesicles biogenesis, some cytoplasmic proteins and also proteins from intracellular organelles (e.g. mitochondria, lysosomes, endosomes) (supplementary table 1). In fact, it was possible to detect the presence of some mitochondrial proteins, such as VDAC-1, ATP synthase F1 subunit beta and subunit alpha within EVs. Heatmap analysis based on the relative abundance revealed that significantly different expressed proteins in NSC EVs are involved in metabolism, cell adhesion and membrane traffic (Figure S4a). The most enriched molecular functions included protein binding, structural molecule activity or catalytic and GTPase activity. In addition, protein–protein interaction network analysis (STRING database) showed that mitochondrial proteins, including ATP synthase subunits in the EVs were interrelated to each other (Figure S4b, c).

**Figure 3.**
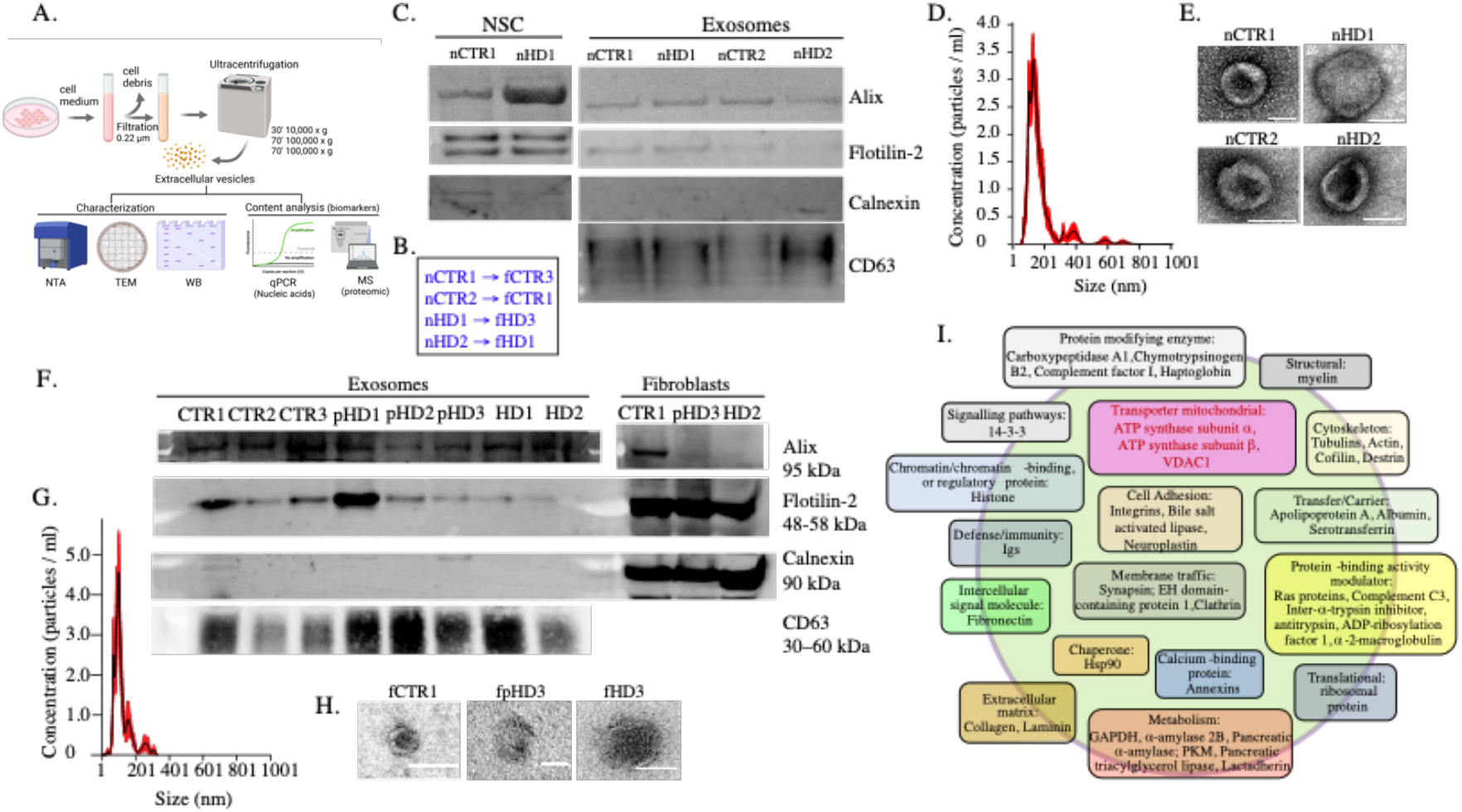
Characterization of EVs isolated from fibroblasts and NSC of HD patients and controls. (a) Schematics of the ultracentrifugation protocol to obtain EVs and further application. (b) Correspondence between differentiated NSC lines and patients’ fibroblasts. (c, f) Western blot of NSC-and fibroblasts-EVs protein markers (Alix, Flotillin-2 and CD63) and Calnexin as a negative control. (d,g) Representative analysis of NSC-and fibroblasts-EVs concentration (particles/ml) and size (nm) obtained by NTA, respectively. (e,h) TEM images of NSC-and fibroblasts-EVs, respectively (scale bar=100 nm). (i) Mitochondrial associated proteins identified by mass spectrometry in NSC-EVs.

The rate of EVs secretion is a cell-specific process dependent on the physiological state or exposure to stimuli (Fader et al., 2008; Ma et al., 2018; Miranda et al., 2018). To study the effect of disease progression on EVs biodistribution, we next examined if the HD cells secreted more EVs. To test this, we evaluated the number of EVs secreted in conditioned media and observed that in HD-fibroblasts and -NSC released more EVs compared to CTRs (Figure 4a, d). Next, we assessed if this increased concentration could be affected by disease-specific parameters. Fibroblasts from older participants and with longer CAG repetitions seemed to release a higher number of EVs suggesting that EVs secretion was increased with disease severity (Figure 4b, c). Importantly, fibroblasts transduced with CD63-GFP or CD63-DsRed, when co-cultured for 24 hours in low density, exchanged single vesicles, as shown in the visualization of green or red puncta in cells with a different CD63 tag (Figure 4e).

**Figure 4.**
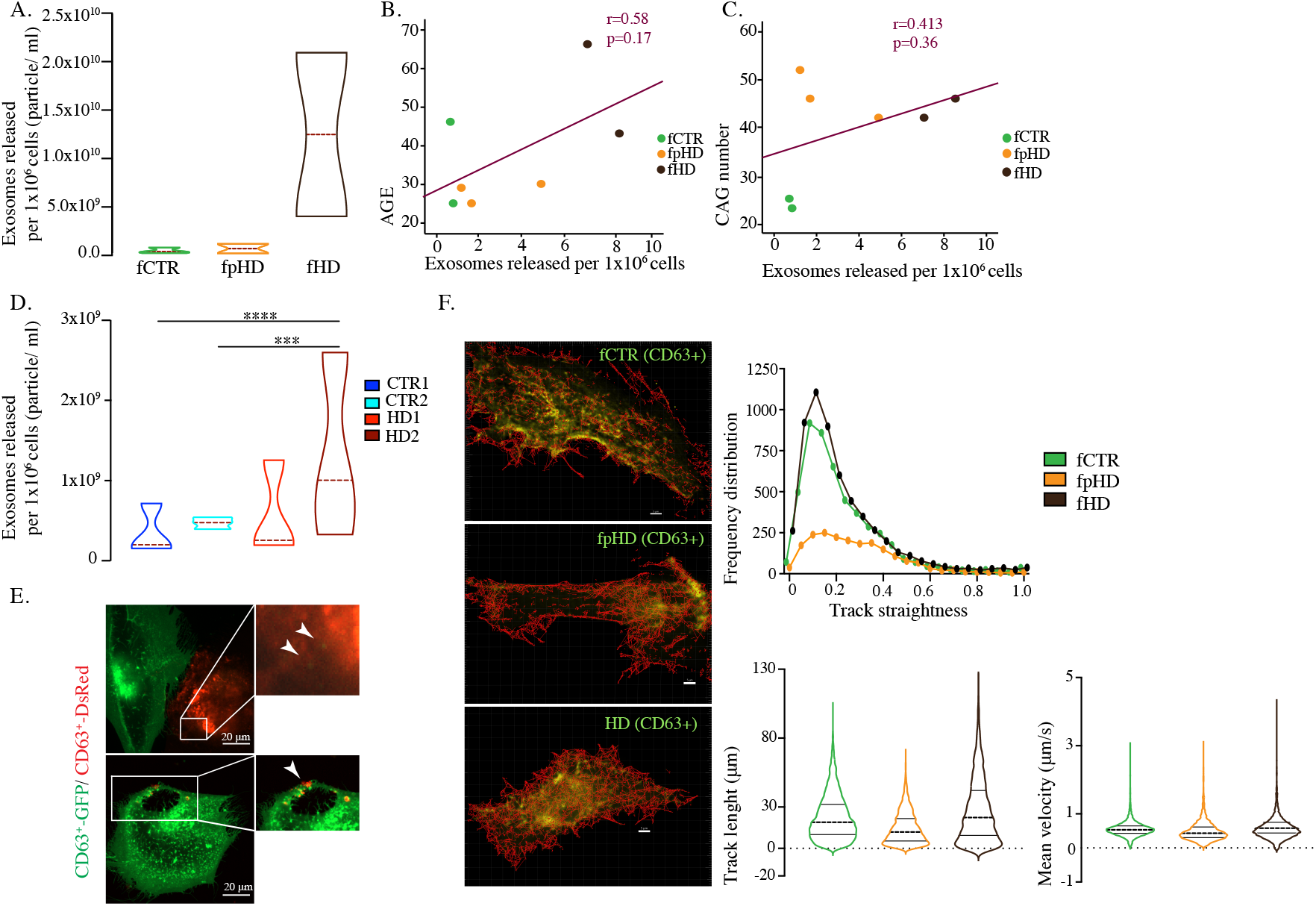
Release of EVs is increased in HD cells. (a) Concentration of EVs released per 1*x*10^6^ fibroblasts calculated by NTA (2 control, 3 premanifest and 2 manifest HD fibroblasts were grouped). Correlation between EVs released per 1*x*10^6^ fibroblasts and patients (b) age and (c) CAG number. (d) Concentration of EVs released per 1*x*10^6^ NSC calculated by NTA. (e) EVs exchange and uptake between fibroblasts transduced with CD63-GFP or CD63-DsRed (scale bar=20 μm). (f) Single particle tracking of exosome dynamics by confocal live cell imaging of CD63-GFP plasma membrane labeled fibroblasts (scale= 5 μm) and corresponding graphics of track straightness, track length (μm) and mean velocity (μm/s). Violin plots represent median with interquartile range. Statistical analysis: one-way ANOVA followed by Bonferroni multiple comparisons test: *** p< 0.001, **** p<0.0001.

The single-particle tracking analysis of CD63+ EVs dynamics (Supplementary videos) shows that trajectories and speed in fpHD and fHD have a similar profile as CTRs (Figure 4f). When evaluating the track straightness of EVs, all cell lines were more prone to random diffusion. Curiously, fpHD presented a decreased frequency distribution.

### HD-extracellular vesicles transport higher levels of mitochondrial genetic material

Mitochondrial DNA copy number is a biomarker of mitochondrial function (Castellani et al., 2020). Since our data shows decreased mitochondrial oxygen consumption rates and altered mitochondrial dynamics in fibroblasts and NSC of HD patients, we investigated the presence of mitochondrial genome and proteins in EVs. Using multiple approaches including long-range PCR, to screen the complete mitochondrial genome, and absolute DNA copy number, we detected mtDNA inside DNAse I-treated EVs (Figure 5a,b,c). Fibroblasts of CTRs and manifest patients have similar mtDNA copy numbers, but the EVs isolated from manifest patients show more mtDNA copies (Figure 5d). Premanifest patients’ cells showed increased mtDNA copy number that was not translated into a corresponding higher EVs mtDNA content (Figure 5e). It is noticeable that HD-EVs transport a higher number of mtDNA copies compared to CTRs-and premanifest-EVs.

The mtDNA copy number was directly correlated with the CAG expansion and number of EVs secreted (Figure 5f). Overall, these data indicate that mtDNA found inside EVs can be a biomarker for disease severity and possibly the generation of mitochondrial-derived vesicles (MDVs) could be a mechanism to counteract defective mitochondrial quality control. Surprisingly, the same results were not observed in NSC. The EVs released from the nHD2 line showed a tendency to have a higher number of mtDNA copies in EVs when compared to control cells. Still, EVs from nHD1 exhibit a very low number of mtDNA copies (Figure 5g).

**Figure 5.**
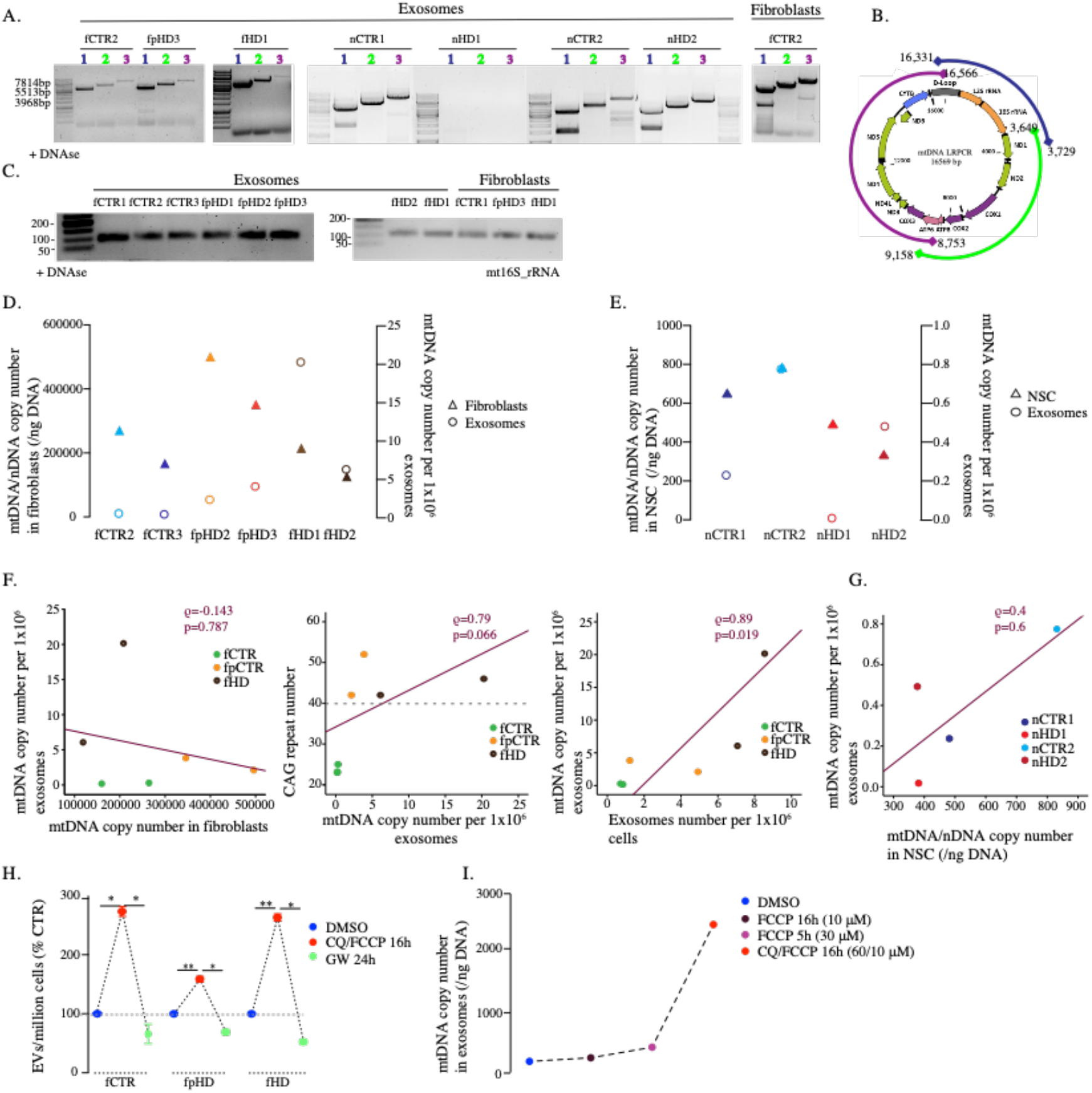
Mitochondrial genome identified in EVs from HD and control cells. (a) Gel electrophoresis images of long-range PCR for mtDNA from purified EVs of fibroblasts (fCTR2-control, fpHD3-premanifest, fHD1-manifest HD lines) and NSC Fibroblast cell line used as controls. (b) schematic of primers for long-range PCR encoding the 16 kbp-size human mitochondrial genome. (c) Representative PCR gel electrophoresis of mtDNA (mt16S) of purified EVs of fibroblasts and NSC. EVs were pre-treated with 10 μg/μl of DNAse I. Fibroblasts cell line used as controls. (d) Absolut mtDNA copy number from fibroblasts and (e) NSC, and corresponding purified EVs. (f) Correlation between mtDNA copy number per 1*x*10^6^ EVs and mtDNA copy number in corresponding cells, CAG number and number of released EVs per 1*x*10^6^ cells, from left to right. (g) Correlation between absolut mtDNA copy number per 1*x*10^6^ EVs and mtDNA/nuclear DNA copy number in NSC, normalized to ng of DNA. (h) Released EVs per 1*x*10^6^ cells treated with chloroquine (CQ) (60 μM) and FCCP (10 μM) for 16h to block lysosomal acidification and promote mitophagy, and GW4869 (30 μM) for 24 h to inhibit the exosomal release. DMSO was used as control. (i) mtDNA copy number in EVs isolated from control fibroblasts treated with FCCP (10 μM) for 16h, FCCP (30 μM) for 5h, and CQ/FCCP (60 μM, 10 μM respectively) for 16h, normalized to ng of DNA. DMSO was used in control cells. Statistical analysis: one-way ANOVA followed by Bonferroni multiple comparisons test: * p< 0.5, ** p<0.01.

Several lines of evidence suggest that there is close coordination between autophagy and the regulation of biogenesis and exosomes released by the cells (reviewed in Xu et al., 2018). Given our observation that cells from manifest patients showed an increase of EVs release, we hypothesized that this is a consequence of autophagy impairment and mitochondrial dysfunction in the late stages of HD. To test this, EVs were isolated from fibroblasts treated with FCCP alone for 5h with three times the concentration of the 24 h incubation (10 μM) and FCCP and CQ simultaneously for 16 hours to induce mitochondrial fragmentation and inhibit the last stage of autophagy, respectively. Treated cells released more EVs than non-treated cells with higher number of mtDNA copies (Figure 5h, i). The FCCP alone induces a discrete increase in a dose-dependent manner suggesting that the mitochondrial-lysosomes axis deregulation is important for the increase shuttle of mtDNA in EVs. Treatment of these cells with GW4869, an inhibitor of EVs release, for 24 hours was used as a negative control. Therefore, the results support our hypothesis that alterations in autophagy efficiency and accumulation of damaged mitochondria could induce the release of EVs with higher content on mtDNA and mitochondrial proteins in HD.

Since a crosstalk between mitochondria and the EVs biogenesis pathways has been suggested (Su Chul Jang et al., 2019) we next performed live-cell microscopy analysis of fibroblasts expressing CD63-GFP or mito-DsRed. After co-cultivation of both cell populations, we observed the exchange of single vesicles (marked in green) and mitochondria (marked in red), as shown in green or red puncta visualization in cells with a different tag (Figure 6a). Moreover, we observed that EVs were able to travel along the lamellipodia toward the nucleus and colocalize with mitochondria (Figure 6b). We further assessed the presence of other mitochondrial proteins within EVs through immunogold-TEM analysis and identified the presence of TFAM, a mitochondrial transcription factor (Figure 6c).

**Figure 6.**
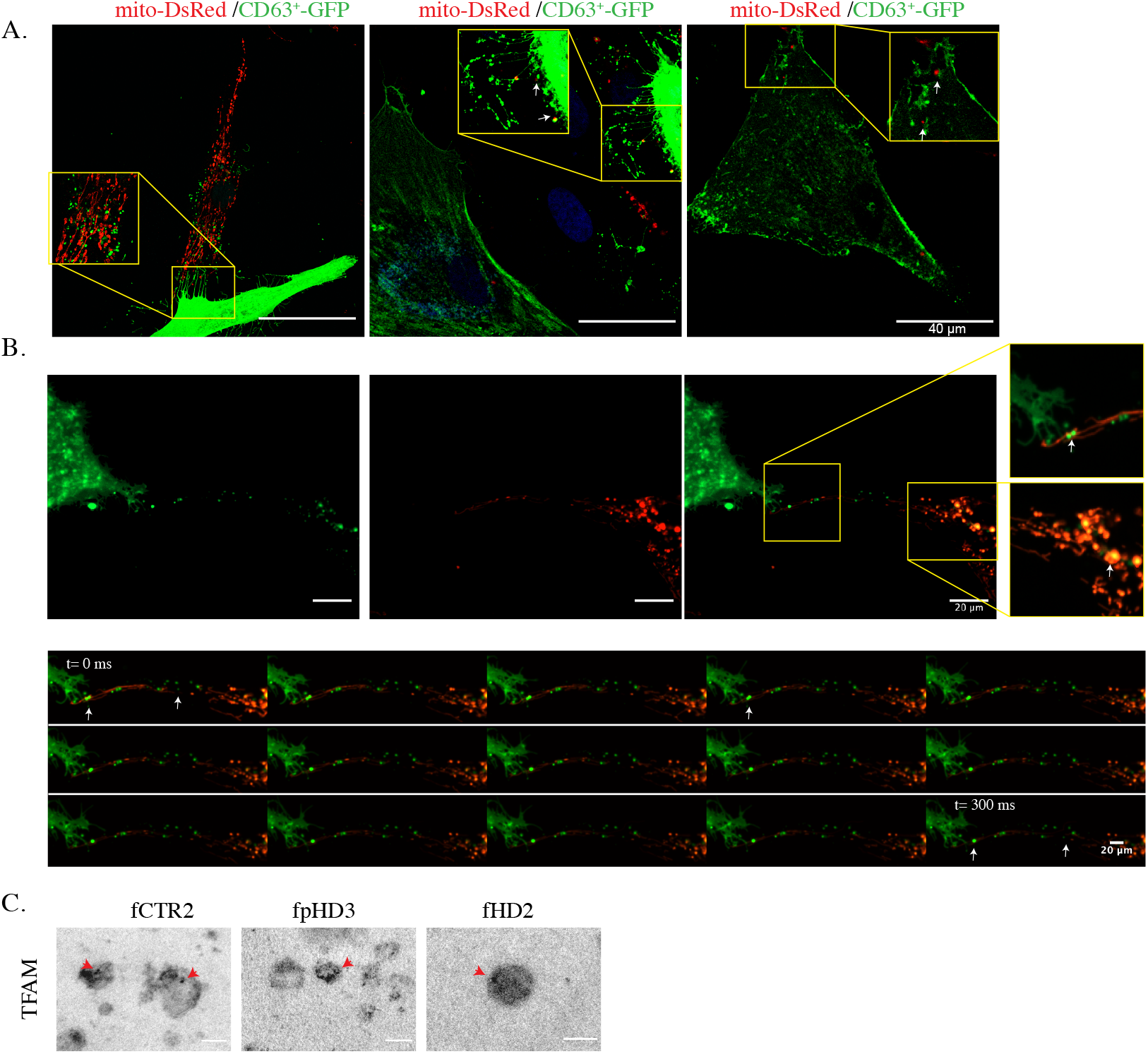
EVs colocalize with mitochondria in fibroblasts. (a) EVs and mitochondria are interchanged between fibroblasts transduced with mito-DsRed or CD63-GFP (scale bar=40 μm). (b) (Upper panel) Colocalization of mitochondria and CD63+-EVs shown by yellow puncta (arrow) (scale bar=20 μm). (Lower panel) Time lapse for 300 ms of exosomal release (arrow) and mitochondrial dynamics in fibroblasts transduced with CD63-GFP and mito-DsRed, respectively (scale bar=20 μm). (c) TFAM Immunogold staining in fibroblasts-EVs (fCTR2-control, fpHD3-premanifest, fHD2-manifest HD lines) (scale bar=100 nm).

### Neuronal-derived EVs from human plasma carry mitochondrial DNA

To validate the potential of mtDNA as a biomarker for HD, total and NDE were isolated from plasma of HD patients and CTRs. NDE are a more specific subpopulation of EVs released by neuronal cells which reflects more accurately the biological mechanisms in neurons than the total pool of EVs present in human plasma. Western blotting analysis confirmed the presence of CD63 in both total EVs and NDE fractions. Moreover, L1CAM (protein of molecular adhesion in neurons) and synaptophysin (marker of presynaptic vesicles) were also detected as neuronal markers in NDE (Figure 7a). Characterization of total EVs and NDE by NTA confirmed the expected size for these vesicles (<200 nm) (Figure 7b). NDE concentration isolated from total human plasma EVs was approximately 5%, as previously described (Sun et al., 2017) (Figure 7c). It was possible to detect the presence of the mitochondrially encoded 16S rRNA gene in the NDE population (Figure 7d). Interestingly, we found that mtDNA copy number was reduced in premanifest patients (30 and 40 copies) compared to prodromal and manifest patients (135 and 157 copies) (Figure 7d), suggesting that the number of mtDNA copies could be a neuronal biomarker for the disease. These data would need to be confirmed in a larger sample size of HD patients and age-matched CTRs to validate its relevance at clinical level.

**Figure 7.**
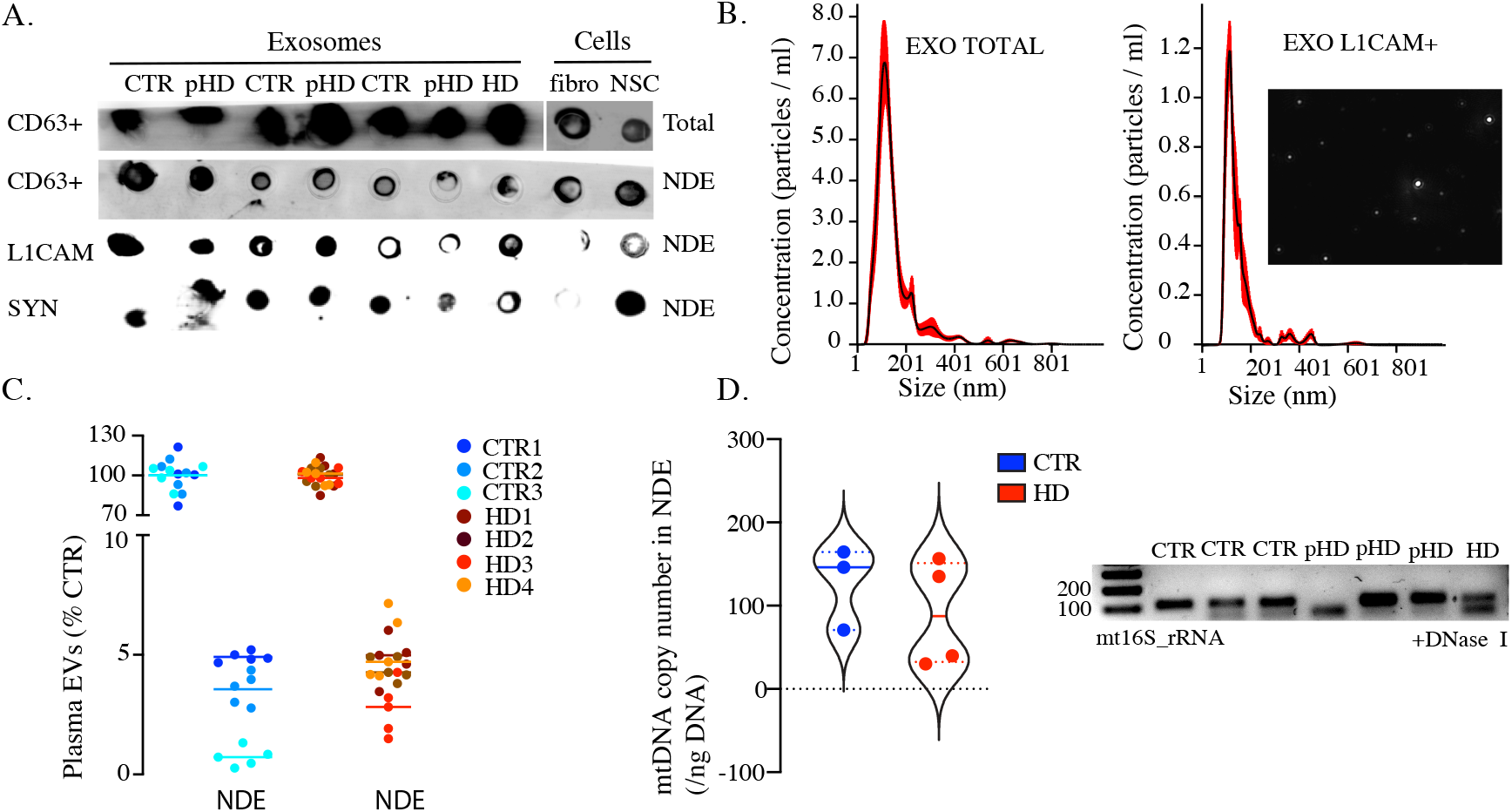
Identification and quantification of neuronal derived exosomes (NDE). (a) Western blot showing NDE protein markers CD63, L1CAM and synaptophysin. (b) Concentration and diameter of plasma exosomes and NDE analyzed by NTA. (c) Percentage of NDE extracted from whole plasma exosomes of HD patients and controls. (D) mtDNA absolute copy number detected in NDE and representative PCR gel electrophoresis. Free DNA was eliminated by DNase I digestion before isolation. Scatter dot plots represent mean±SEM and violin plots median with interquartile range.

## DISCUSSION

Disruption of mitochondrial function and autophagy mechanisms are well known features of HD (Lopes, 2022; Lopes, 2020; Rui et al., 2015). An intricate communication exists between mitochondrial dysfunction and cellular degradation pathways in neurodegenerative disorders.

Here, we demonstrate that NSC differentiated from HD patient’s derived iPSC exhibited features of mitochondrial and metabolic impairment associated with reduced levels of PGC-1α, the master regulator of mitochondrial biogenesis. We previously documented in fibroblasts from the same donors corresponding to patients with manifest HD a general decrease in mitochondrial bioenergetic profile, increased levels of mitochondrial ROS production, and fragmentation associated with reduced mtDNA copy number compared with premanifest HD patients (Lopes, 2022). Because defective mitochondria are eliminated by autophagy/mitophagy, we further explored in this study the impairment of autophagosomal pathway in these fibroblasts. We found that the levels of autophagy/mitophagy protein markers are increased in cells from manifest HD, suggesting an impaired autophagic degradation and/or increased autophagic induction. Of note, the accumulation of LAMP2, p62 and LC3-II in HD cells supports the hypothesis of impaired autophagosome/lysosome fusion once p62 is degraded via autophagy and the protein levels are inversely related to autophagic activity, LC3-II is proportional to the number of autophagosomes and LAMP2 protein is related to the levels of lysosomes (Yoshii & Mizushima, 2017). Disruption of mitochondria function can inhibit lysosomal activity and trigger the accumulation of large vacuoles positive for lysosomal markers in a mechanism mediated by ROS (Demers-Lamarche et al., 2016). Mitochondria are highly dynamic membraned organelles dependent on efficient mitochondrial quality control (MQC) to ensure correct mitochondrial plasticity, disposal, and replenishment mandatory for cellular homeostasis. Several pathways are involved in mitochondrial MQC including mitophagy and mitochondrial-derived vesicles (MDVs) generation. Mitophagy is a multi-step process where the dysfunctional organelle is initially engulfed into an autophagosome, which later fuses and delivers the cargo to lysosomes for degradation. Recently mitochondrial-lysosomal membrane contact sites were suggested to modulate mitochondrial dynamics either through intracellular calcium regulation or by regulating mitochondrial fission (Peng et al., 2020; Wong et al., 2018). Another line of defense involved in the elimination of damaged, oxidized mitochondria independent of mitochondrial depolarization, autophagy signaling, or mitochondrial fission are MDVs (Cadete et al., 2016; Neuspiel et al., 2008). Initially it was proposed that the generation and upload of MDVs was regulated through a mechanism dependent on PINK1 and Parkin (McLelland et al., 2014). Recently a new MDVs biogenesis pathway dependent on MIROs and DRP1 was described. Moreover, MDVs transport not only single proteins but also can transport fully assembled complexes to lysosomes (König et al., 2021). The MDVs cargo can have two fates, the incorporation into the endolysosomal system through the fusion with multivesicular bodies (MVBs), followed by lysosomal degradation or secretion as EVs, or can be deliver into peroxisomes (D’Acunzo et al., 2021; Matheoud et al., 2016; Neuspiel et al., 2008; Soubannier et al., 2012). Several subtypes of MDVs have been identified based on their cargo selectivity and destination. These include the mitochondria-anchored protein ligase – containing MDVs, the pyruvate dehydrogenase (PDH)+/TOM20 - which traffics oxidized molecules into lysosomes and TOM20+/PDH-MDVs that also targets the lysosomes but is a PINK1/Parkin independent pathway (McLelland et al., 2014; Soubannier et al., 2012).

Mitovesicles are another subclass of MDVs of endosomal origin that are secreted as EVs but are enriched in mitochondrial proteins such as VDAC, COX-IV and PDH-E1α (D’Acunzo et al., 2021). The proteomic analysis surprisingly revealed the presence of almost all electron transport chain and the PDH complex subunits (D’Acunzo et al., 2021). In this sense lysosomal dysfunction and associated impaired autophagy can promote the release of MDVs through the endosomal pathway. Our data shows that manifest HD cells display mitochondrial dysfunction, increased mitophagy and impaired lysosome-mediated autophagy associated with an increased release of EVs and higher number of mtDNA copies. Interestingly when we reproduced these conditions in control cells by inducing mitophagy and inhibiting autophagy the mtDNA content on EVs increased dramatically. Moreover, mutant HTT sequestration of autophagy proteins could exacerbate the disease progression by an age-dependent manner and EVs secretion could be a compensatory mechanism to eliminate cellular stressors (Lee et al., 2012). Adipocyte cells also responded to mitochondrial stress and inhibition of lysosomal degradation by rapidly and robustly releasing sEVs that contain oxidized mitochondria (Crewe et al., 2021). Interestingly, when these EVs are accumulated by cardiomyocytes, they can induce a transient mitochondrial oxidative stress, resulting in a protective compensatory antioxidant signaling (Crewe et al., 2021). In addition, as previously shown by other reports in breast cancer cells, fibroblast-derived EVs colocalize with mitochondria indicating that EVs can possibly fuse with targeted mitochondria in cells (P. Sansone et al., 2017). Our proteomic analysis of the EVs content validated previously established mitochondrial protein content (S. C. Jang et al., 2019; Peruzzotti-Jametti et al., 2021; P. Sansone et al., 2017). In general, the proportion of mitochondrial proteins described to be secreted within EVs is up to 10% of the total proteome (Sugiura et al., 2014). Increased EVs-associated secretion of mitochondrial proteins and DNA to compensate for the lysosomal impaired degradation add a new mechanism to the paradigms of MQC. Mito/autophagy are the classical pathways involved in removing of dysfunctional mitochondria, but when this degradative process is compromised, MDVs formation most likely complements this process by shuttling damaged, oxidized cargo through the endosome/exosome secretory pathway. Furthermore, recent evidence suggests that EVs can transfer between NSC structurally and functionally intact mitochondria with the ability to recover mitochondrial function in mtDNA-depleted cells paving new ways to the development of treatment strategies in neurodegenerative disorders (Peruzzotti-Jametti et al., 2021). In addition, we also detected EVs-associated mtDNA in plasma-derived NDE from HD patients reinforcing the hypothesis of using NDE mtDNA as a biomarker for monitoring disease progression. EVs derived from neurons are a new and controversial field that holds great potential as a liquid biopsy for the diagnostics of central nervous system diseases. The studies that addressed the NDE cargo focused mainly on microRNA or protein of interest for the studied disease and only a few have concentrated on mitochondrial proteins (Goetzl et al., 2021).

The role of MDVs as an MQC-associated mechanism or even its implication on the neuropathological processes underlying neurodegenerative disorders is unclear. MtDNA is commonly found in EVs secreted from different cells, yet the biological function is largely unknown (Elzanowska et al., 2021). Additionally, a decline in mtDNA levels with age in EVs from human plasma was reported (Lazo et al., 2021), supporting our hypothesis that the increased mtDNA levels in HD older patients that we found is disease-dependent.

The mitochondrial content found in EVs, particularly mtDNA, is increasingly recognized as an agonist of the innate immune system capable of inducing an inflammatory response through various pattern-recognition receptors, including Toll-like receptors, NOD-like receptors and interferon-stimulatory DNA receptors (West & Shadel, 2017). Pro-inflammatory mitochondrial proteins loading into EVs capable of induce an immune response by triggering microglia to release pro-inflammatory factors to the extracellular media and consequently promote cell death have been reported in neurodegenerative diseases (Picca et al., 2019; K. Todkar et al., 2021; X. Wang et al., 2020).

We understand that every work has room for improvement and, like so, this study has some limitations: 1) the number of participants and the disease stage representativity. A comprehensive study enrolling more HD patients and age-matched controls is necessary to better understand the impact of these results in the disease pathophysiology; 2) these results should be further confirmed in EVs released from patient’s iPSC-derived neurons. However, several questions remain to be answered concerning EVs roles. Are EVs a mechanism to encapsulate and transfer mitochondrial material, genetic or proteic, to recipient cells, aiming to rescue their mitochondrial function, or EVs-mediated secretion is an alternative/compensatory mechanism, once autophagy is impaired, to eliminate defective mitochondria.

In conclusion, our data provide evidence of an innovative mechanism involving a dysfunction on the mitochondrial-lysosomes axis that promotes the release of damaged mitochondrial components within EVs. The MQC process comprises several pathways and, significantly in HD, the MDVs trafficking seems to be part of a degradative mechanism complementary to mitophagy.

## ABBREVIATIONS

ALS: amyotrophic lateral sclerosis
BDNF: brain-derived neurotrophic factor
BSA: bovine serum albumin
CES: collision energy spread
CNS: central nervous system
CQ: chloroquine
CTRs: controls
CUR: curtain gas
DAMP: damage-associated molecular patterns
DOG: 2-deoxyglucose
ECAR: extracellular acidification ratio
EVs: extracellular vesicles
FBS: fetal bovine serum
FDR: False Discovery Rate
HBSS: Hanks’s Balanced Salt solution
HD: Huntington’s disease
HTT: huntingtin
ID: internal diameter
IDA: information-dependent acquisition
iPSC: induced-pluripotent stem cells
MQC: mitochondrial quality control
mtDNA: mitochondrial DNA
MDVs: mitochondrial-derived vesicles
MVBs: multivesicular bodies
NDE: neuronal-derived EVs
NSC: neural stem cells
NTA: Nanosight Tracking Analysis
OCR: oxygen consumption rate
PDH: pyruvate dehydrogenase
PVDF: polyvinylidene difluoride
qPCR: real-time PCR
ROS: reactive oxygen species
RT: room temperature
sEVs: small EVs
TEM: transmission electron microscopy.

## ACKNOWLEDGEMENTS

We acknowledge Dr. Sandra Mota and Dr. Paulo Oliveira for their critical review of the manuscript. Dr. Mónica Zuzarte, for fibroblasts and NSC TEM analysis at the ‘Laboratório de Bio-imagem de Alta Resolução’ of the Faculty of Medicine of the University of Coimbra; Dr. Teresa Rodrigues and Dr. Henrique Girão from Institute of Biomedical Imaging and Life Sciences (IBILI) of the Faculty of Medicine of the University of Coimbra for facilitating the access to NTA equipment Lab; Nurse Margarida from Neurology service of the ‘Centro Hospitalar da Universidade de Coimbra’ for the liaison with patients.

## ETHICAL APPROVAL AND CONSENT TO PARTICIPATE

Ethical approval was given by the Medical Ethics Committee of Coimbra Hospital and University Centre (CHUC).

## AVAILABILITY OF SUPPORTING DATA

The supporting data and materials are provided in Supplementary Figures and Tables.

## Author Contributions

CL designed this study, MB, RV, CL, SA performed the experiments, MB, RV, CL analyzed the data, MB, CL drafted the initial manuscript, MB, RV, CL, CR revised this manuscript; all authors read and approved the final manuscript.

## Supplementary Data

**Figure S1.**
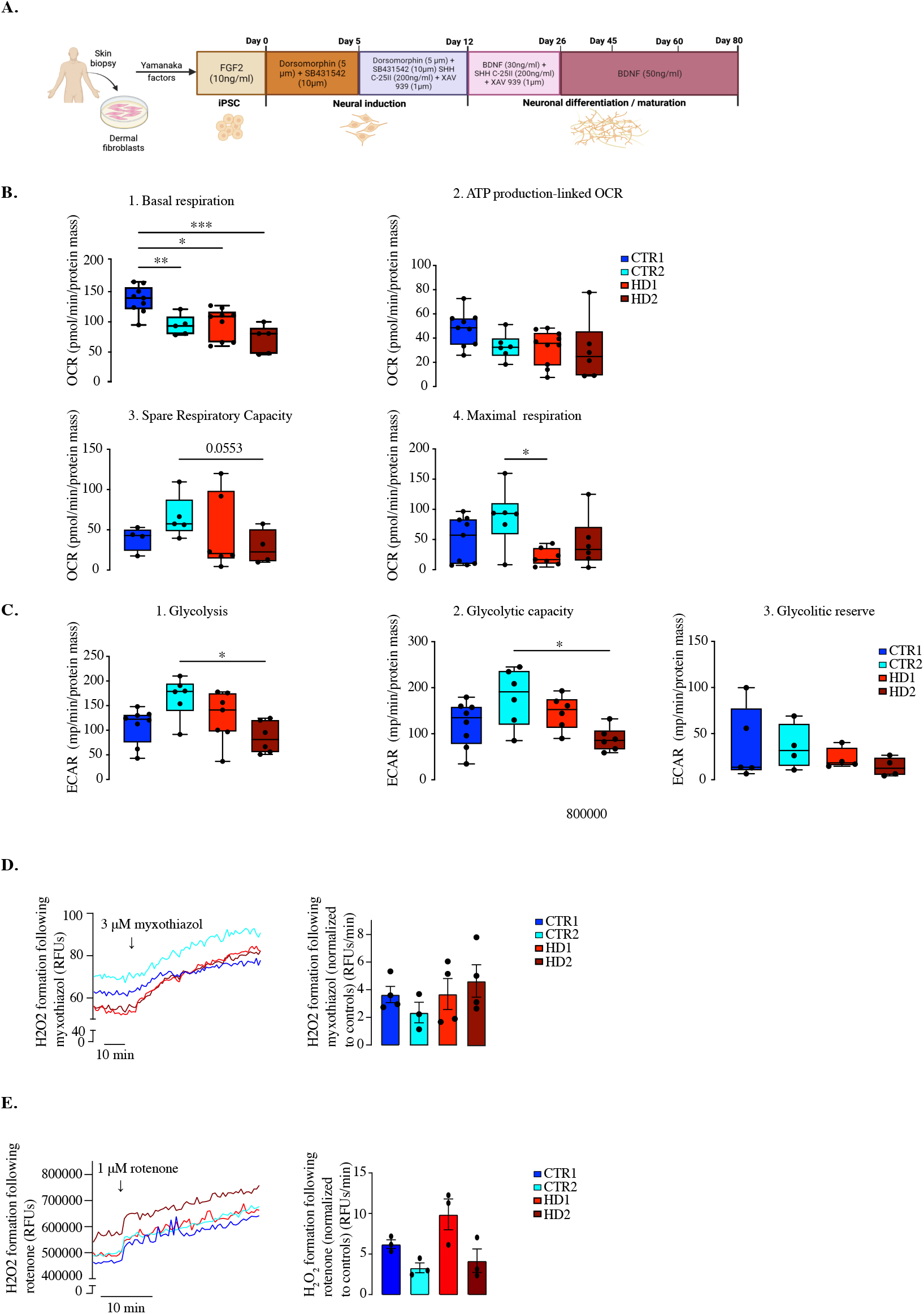
Characterization of neuronal cells differentiation protocol, mitochondrial bioenergetics and hydrogen peroxide formation. (a) Differentiation protocol to obtain striatal-like neurons from iPSC reprogrammed of human fibroblasts. Timeline for morphogens/factors and concentrations are indicated. To obtain proliferative NSC the differentiation protocol is only applied until day 12-15 and no SHH C-25 II is added. (b) Analysis of oxygen consumption rate (OCR) parameters in NSC from HD patients and CTRS: basal respiration, ATP production, spare respiratory capacity and maximal respiration (N=5-10). (c) Analysis of extracellular acidification rate (ECAR) in NSC from HD patients and controls: glycolysis, glycolytic capacity and glycolytic reserve (N=5-10). (Left) (d) Basal mitochondrial hydrogen peroxide production was measured using MitoPY1 fluorescence (RFUs) for 10 min and after incubation with myxothiazol (3 μM) and (e) rotenone (1 μM) (inhibitors of mitochondrial complexes III and I, respectively) for 20 min. (Right) Bar plots represent the rate of hydrogen peroxide formation after stimulus (RFUs/min) normalized to controls (N=3-4). Box and whiskers represent median and data distribution, bar and scatter dot plots represent mean±S.E.M and violin plots represent median with interquartile ranges. Statistical analysis: one-way ANOVA followed by Bonferroni multiple comparisons test: * p<0.05, ** p<0.01, *** p< 0.001.

**Figure S2.**
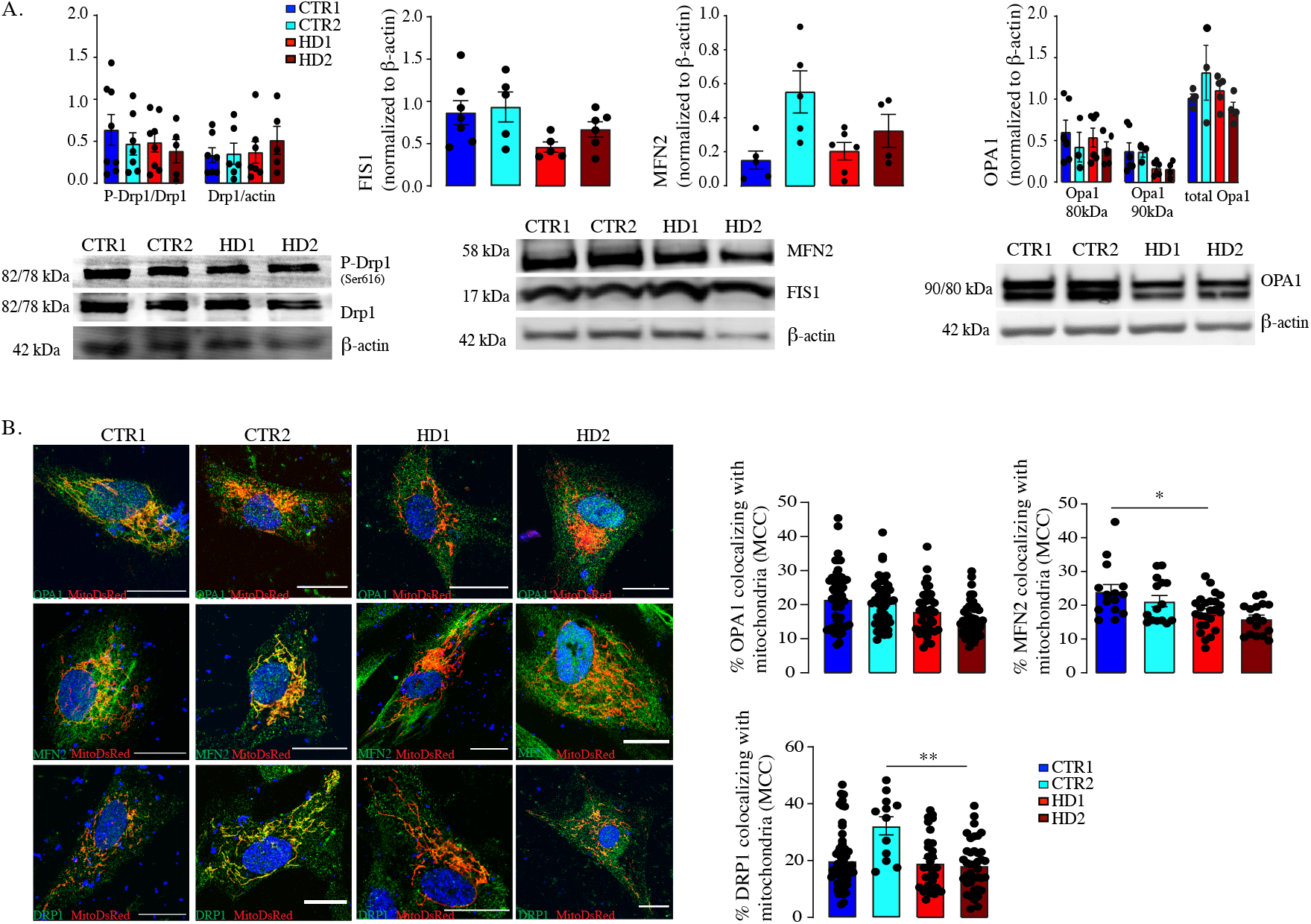
Western blot and immunofluorescence analysis of mitochondrial proteins OPA1, MFN2 (fusion) and DRP1, FIS1 (fission) molecules in pDsRed2-Mito transfected NSC. (a) Western blot analysis of mitochondrial fission-(p-Drp1 (Ser616), Drp1 and FIS1) and fusion-associated (MFN2 and OPA1) proteins in NSC from HD patients and CTRS. Results were normalized to ß-actin levels (N=3-8). (b) (Right) Colocalization represented by Mander’s coefficient of colocalization (MCC) of fusion/fission protein and mitochondria in NSC transfected with pDsRed2-Mito. (Left) Representative images were photographed at ×63 (scale bar=20 μm) and colocalization quantification was performed on Z-stacks using the JACoP ImageJ plugin. Bar plots represent mean±S.E.M (N=12-57). Statistical analysis: one-way ANOVA followed by Bonferroni multiple comparisons test: * p< 0.05, ** p<0.01.

**Figure S3.**
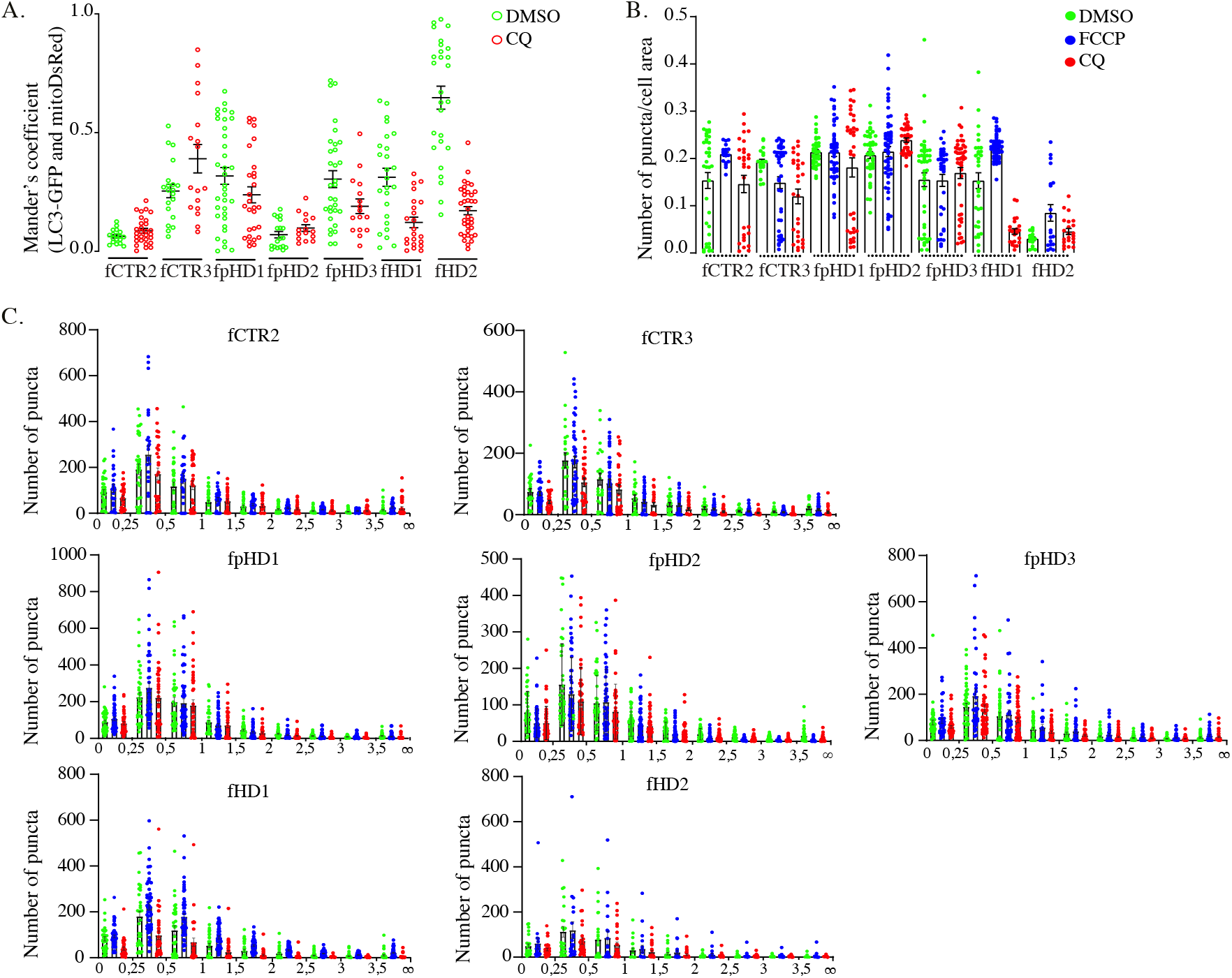
Effects of autophagy modulators on lysosomal colocalization with mitochondria. (a) Colocalization, represented by Mander’s coefficient, of LC3 and mitochondria in fibroblasts transduced with mitoDsRed and GFP-LC3 and treated with CQ (100 μM) for 5h (N>=17). (b) Puncta number analyzed per cell area in cells treated with CQ (100 μM) and FCCP (30 μM) for 5h. (c) A histogram representing the frequency distribution of the estimated sizes of LC3 puncta in fibroblasts treated with CQ (100 μM) or FCCP (30 μM) for 5h. The data was sorted into bin-size and a frequency distribution histogram was plotted. Area is represented in μm^2^ (XX axis) (N>=20). Bar and scatter dot plots represent mean±S.E.M..

**Figure S4.**
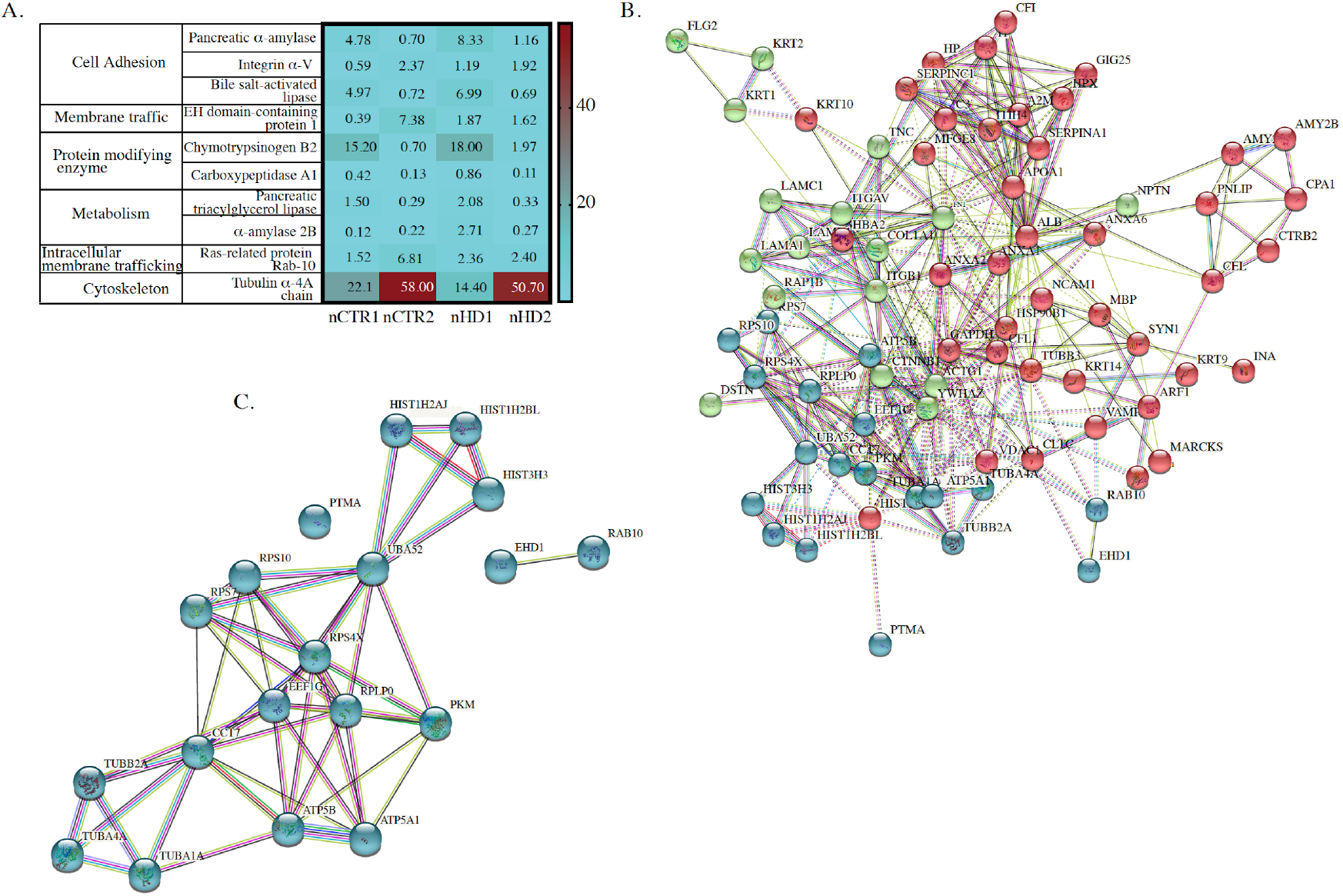
Heatmap of gene ontology (GO) enrichment analysis of differentially expressed proteins generated by mass spectrometry analysis in control and HD NSC lines. (a) GO protein class enrichment analyses of up-regulated and down-regulated proteins were performed using The PANTHER Classification System and (b) the protein-protein interaction network and identification count were obtained from the STRING database.

**Video 1, Video 2 and Video 3 shows intracellular exosome movement in fibroblasts expressing CD63-GFP protein.**

The movement of CD63-GFP+ in human fibroblasts was monitored for 5 min with a frame rate of 1.5 s/frame by live cell confocal. Each video represents a cell line: 1-CTR, 2 – premanifest HD, 3 -manifest HD.

**Supplementary Table 1.**
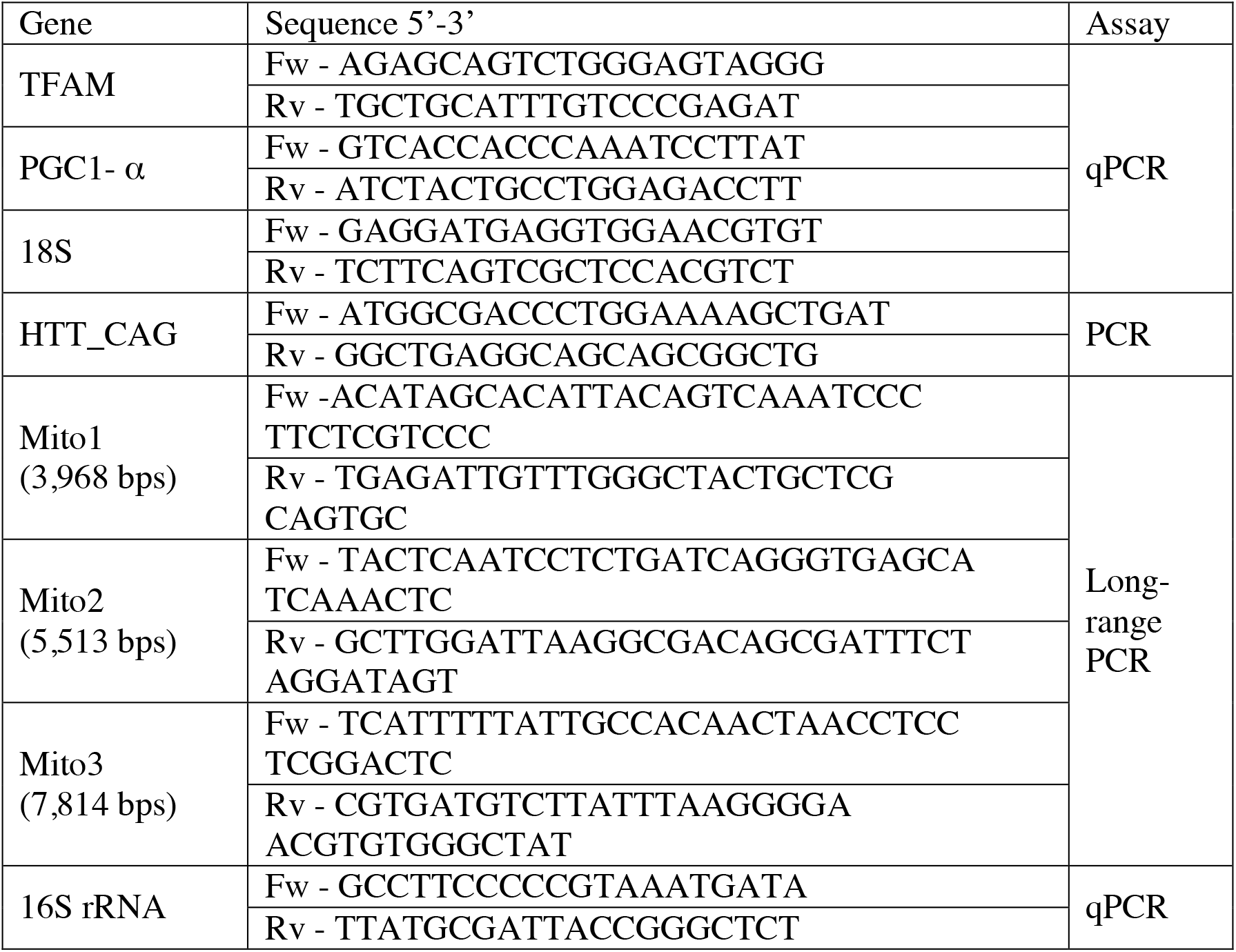

## Supplementary Materials and Methods

### SWATH-MS analysis

The chromatographic separation was performed on a YMC-Triart C18 Capillary Column 1/32′′ (12 nm, S-3 μm, 150 mm × 0.3 mm) and using a YMC-Triart C18 Capillary Guard Column (0.5 μm × 5 mm, 3 μm, 12 nm) at 50. The flow rate was set to 5 μLmin^-1^ and mobile phases A and B were 5% DMSO plus 0.1% formic acid in water or acetonitrile, respectively. The LC program was performed as followed: 2-5% of B (0–2 minutes), 5– 28% of B (2–50 minutes), 28–35% of B (50–51 minutes), 35–98% of B (51–52 minutes), 98% of B (52–61 min), 90–2% of B (61–62 min), and 2% of B (62–68 min). The ionization source was operated in the positive mode set to an ion spray voltage of 5,500 V, 25 psi for nebulizer gas 1 (GS1) and 25 psi for the curtain gas (CUR).

Quantitative data processing was conducted using SWATH™ processing plug-in for PeakView™ (v2.0.01, ABSciex®) (Lambert et al., 2013). After retention time adjustment using the MBP-GFP peptides, up to 15 peptides, with up to five fragments each, were chosen per protein, and quantification was attempted for all proteins in the library file that were identified from ProteinPilot™ search. Only proteins with at least one confidence peptide (FDR<0.01) in no less than four replicate conditions and with at least three transitions were considered. Peak areas of the target fragment ions (transitions) of the retained peptides were extracted across the experiments using an extracted-ion chromatogram (XIC) window of 4 minutes with 100 ppm XIC width.

The proteins’ levels were estimated by summing all the transitions from all the peptides for a given protein that met the criteria described above (an adaptation of Collins et al., 2013) and normalized to the levels of the internal standard (Anjo et al., 2019).

A Kruskal–Wallis test was performed to identify the proteins differentially regulated between all the comparisons followed by the Dunn’s test of Multiple Comparisons, with Benjamini–Hochberg p-value adjustment, to determine in which comparisons statistical differences were observed. All the analyses were performed using the normalized protein levels and a p-value of 0.05 were defined as cut-off.

